# A fly model establishes distinct mechanisms for synthetic CRISPR/Cas9 sex distorters

**DOI:** 10.1101/834630

**Authors:** Barbara Fasulo, Angela Meccariello, Maya Morgan, Carl Borufka, Philippos Aris Papathanos, Nikolai Windbichler

## Abstract

Synthetic sex distorters have recently been developed in the malaria mosquito, relying on endonucleases that target the X-chromosome during spermatogenesis. Although inspired by naturally-occurring traits, it has remained unclear how they function and, given their potential for genetic control, how portable this strategy is across species. We established *Drosophila* models for two distinct mechanisms for CRISPR/Cas9 sex-ratio distortion - “X-shredding” and “X-meddling” - and dissected their target-site requirements and repair dynamics. X-shredding relies on sufficient meiotic activity of the endonuclease to overpower DNA repair and can operate on a single repeat cluster of non-essential sequences. X-meddling by contrast, i.e. targeting putative haplolethal X-linked genes, induced a bias towards males that is coupled to a loss in reproductive output, although a dominant-negative effect may drive the mechanism of female lethality. Our model system will guide the study and the application of sex distorters to medically or agriculturally important insect target species.

## Introduction

In a population of sexually reproducing organisms, a significant sex bias towards males is predicted to decrease the population’s overall reproductive output. This is because the fecundity of females, the sex with a lower rate of gamete production, generally determines the size of a population. Forcing the sex-ratio towards males has thus long been regarded as a potential avenue for the genetic control of harmful insect pest or disease-vector populations. While the introduction of various forms of female-killing genetic traits could achieve this goal, more powerful strategies have also been theorized. Hamilton, for example, speculated that a population of a heterogametic species would become increasingly male-biased if, at each generation a mutant Y chromosome would favour its transmission over the X-chromosome. The decline in female numbers would result in a reduced population size and eventually the collapse of the population (Hamilton 1967). Hamilton’s thinking was inspired by records of distorter traits in *Aedes aegypti* and *Culex pipiens* that can produce extreme sex-ratios of >90% males. Cytological observations during male meiosis showed broken X-chromosomes suggesting a causal link with the male-bias phenotype (Sweeny and Barr 1978, Newton M.E. 1976). These findings inspired the generation of artificial distorter traits (Windbichler, Papathanos, and Crisanti 2008) first by using His-Cys box homing endonucleases and subsequently RNA-guided endonucleases. In the malaria vector *Anopheles gambiae* (*A. gambiae*) autosomal I-PpoI (Galizi et al. 2014) or CRISPR/Cas9-bearing transgenes (Galizi et al. 2016) were used to target sequences on the X-chromosome during male meiosis. Cage experiments using such distorter traits induced extreme male-biased sex-ratios and population collapse confirming the potential of this system for genetic control.

Mechanistically, the nature of the target locus for which successful X-shredding was demonstrated suggested a possible coalescence of different effects in the mosquito system. The target sites are situated within the *Anopheles gambiae* 28S rDNA cluster which simultaneously represents (i) a high-copy number repeat on the X-chromosome, (ii) an essential gene for ribosome biogenesis and function, (iii) the nucleolar organizing region of the cell as well as (iv) a sequence adjacent to the centromere of the X-chromosome and (v) the predicted pseudo-autosomal region of the X-chromosome mediating pairing with the Y chromosome during meiosis (Hall et al. 2015). The possible conflation of effects induced by X-shredding in the mosquito has been a source of uncertainty regarding the potential to transfer this paradigm to other important pest species. In particular, the X-shredding approach has not been clearly delineated from a related, recently-proposed strategy, based on the targeting of X-linked haploinsufficient genes (commonly ribosomal genes) with the intent to induce female lethality (Burt and Deredec 2018). To delineate this mechanism, which is assumed not to alter gamete production and to come into effect only in the developing progeny, we refer to it as X-meddling. X-meddling would also generate a male biased progeny but would be expected to lead to a significant loss of reproductive output in the form of inviable female embryos. Reduced hatching is however a feature of the mosquito sex distortion system in some transgenic strains (Galizi et al. 2014). Although experiments suggested that carry over of endonuclease protein rather than insufficiency of the rDNA was responsible for this zygotic lethality (Windbichler, Papathanos, and Crisanti 2008), a contribution of altered target gene function to this effect could not be ruled out completely.

Here, we have sought to disentangle and reconstitute these two strategies in *Drosophila melanogaster* by targeting with CRISPR/Cas9 both X-linked multicopy repeats and, in parallel, X-linked putative haplolethal genes essential for ribosome function. By doing so we have also sought to demonstrate that the X-shredding mechanism is transferable, in principle, between species and between target genes and has potential applications beyond malaria vector control. We have also explored the efficiency of X-meddling in biasing the sex-ratio as an alternative strategy for genetic control.

## Results

Due to the inadequacies of the Gal4/UAS system for directing expression in later stages of spermatogenesis (White-Cooper 2012), we first generated multiple strains where *cas9* or *cpf1* (*cas12a*) were placed under the direct transcriptional control of the *βtub85Dtub85D* promoter. This regulatory element drives high expression of genes exclusively in the male germline, during the primary spermatocyte stage and, as demonstrated in mosquito (Windbichler, Papathanos, and Crisanti 2008, Galizi et al. 2014), this is the stage with the strongest evidence of X-shredding activity.

To evaluate CRISPR function, we first crossed *βtub85Dtub85D-*endonuclease bearing lines to lines containing gRNAs targeting the X-linked, single copy *white* gene, used here as a phenotypic marker (gRNAs *w_ex3_2* for Cas9 and *w_ex3_1* for Cas12a) (Kondo and Ueda 2013). The mutation of *white* results in individuals with white eyes due to a lack of pigment and we compared the activity of the different lines by counting the fraction of offspring with white eyes (Figure S1). We found variability of mutation frequencies between the different *βtub85Dtub85D-cas9* lines, depending on the chromosomal location of the endonuclease transgene. In addition, the level of activity of all the *βtub85Dtub85D* driven lines was significantly lower when compared to *cas9* driven by the *nanos* (*nos*) promoter using the same gRNA (96.7% white eyes). LbCfp1 showed low levels of activity (0.6% white eyes) and for our further experiments we exclusively utilized Cas9. Unless specifically indicated, we used the *βtub85Dtub85D-cas9*^*20F*^ line, which yielded the highest *white* mutation frequency of 52.2% for all experiments. To identify X-linked sequences which could be targeted by CRISPR/Cas9 in the male germline we used two different approaches. First, using publicly available short read and long read *Drosophila* datasets, we employed the Redkmer pipeline (Papathanos and Windbichler 2018) which we previously developed to identify putative X-linked repeat sequences from raw sequence data alone (Figure S2). Compared to *A. gambiae* (Papathanos and Windbichler 2018) this analysis predicted that the X-chromosome of *Drosophila* harboured less abundant X-chromosome repeats (<250 copies) that were not also present on other chromosomes (Figure S3). From the set of kmers of 25 nucleotides that passed our criteria including abundance, X-linkage, and the lack of predicted off-target cleavage, we selected 8 target sequences for gRNA design located in multiple sites across the X-chromosome (Figure S2, S3). These target repeat sequences were predicted to be confined to single chromosomal regions with the most abundant sequences located within 5 annotated genes (*esi-2.1*, *muc14a*, *hydra*, *CG33235* and *CG15040*). We made no assumption about the higher-order structure or the conservation of these putative repeat sequences between individuals, although all chosen gRNA repeat targets were also predicted to be multicopy sequences on the X when we searched the DmelR6.01 genome assembly (Figure 1B). In a second approach, we identified putative haploinsufficient genes (Marygold et al. 2007), on the *Drosophila* X-chromosome and designed four gRNAs targeting conserved regions within the *RpS5a* (McKim, Dahmus, and Hawley 1996), *RpS6* (Stewart and Denell 1993) ribosomal protein genes.

**Figure 1.**
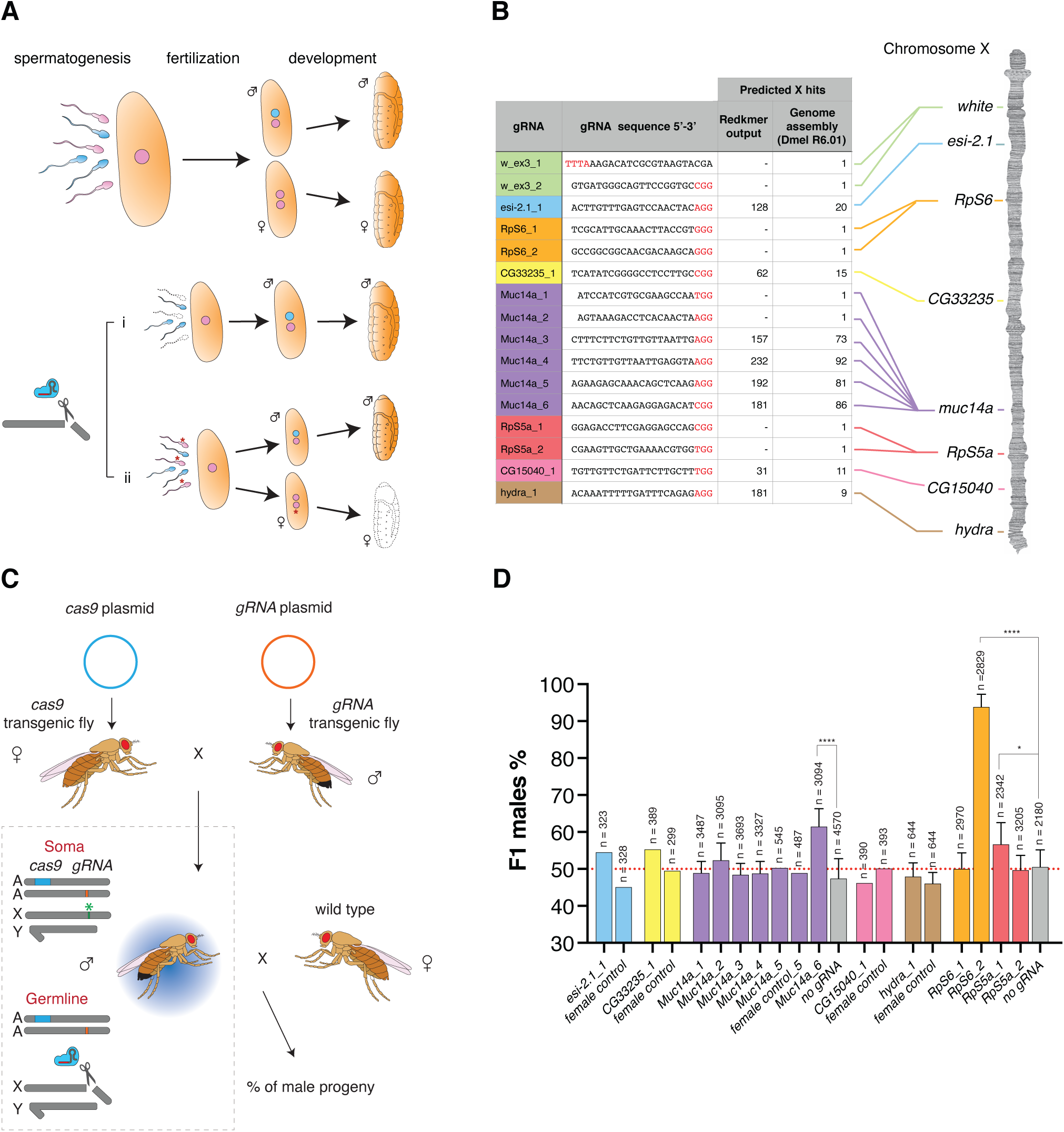
Development of sex ratio distorters in *Drosophila*. **(A)** Model for pre-zygotic and post-zygotic effects on the reproductive sex-ratio. Compared to the unaltered process of development (top), an endonuclease targeting the X-chromosome during spermatogenesis (bottom) could either negatively affect the production or function of X-bearing sperm (i, prezygotic effect) or introduce genetic modifications that are detrimental to females inheriting such modified X-chromosomes during development (ii, postzygotic effect; e.g. the mutation of haploinsufficient genes). Pink: X-chromosome bearing nucleus. Cyan: Y chromosome bearing gamete. The asterisk indicates gametes bearing modifications at the target site. **(B)** gRNA target sequences used in this study and their positions on the X-chromosome. The protospacer-adjacent motif (PAM) is indicated in red, and the X-chromosome hits predicted by Redkmer and the number of predicted perfect BLASTn hits to the X-chromosome in the *Drosophila* genome assembly (Dmel R6.01) are indicated. **(C)** Schematic of the experimental crosses. Females bearing the *cas9* transgene were crossed to males carrying the *gRNA* transgene. Trans-heterozygote males were then crossed to wild-type females. Cas9 in concert with the gRNA cleaves the X-chromosome at the target site (green asterisk) in the germline and the effect is measured as the sex-ratio of the progeny. A: autosome; X: X-chromosome, Y; Y-chromosome. **(D)** Male seX-ratios in the offspring from crosses of *βtub85Dtub85D-cas9/gRNA* with wild-type *w* females. Progeny of *βtub85Dtub85D-cas9/*+ males crossed to wild type females (no gRNA) or from the reverse cross (*βtub85Dtub85D-cas9/gRNA* females crossed to wild type males = female control) served as a control. Crosses were set as pools of males and females or as multiple male single crosses in which case error bars indicate the mean ± SD for a minimum of ten independent single crosses. For all crosses n indicates the total number of individuals (males + females) in the F1 progeny counted. P-values *p < 0.05, ****p < 0.0001.

All gRNAs were cloned downstream of the ubiquitous Pol III promoter of the *Drosophila* U6 snRNA gene and we generated independent gRNA expressing lines for all selected gRNAs. In addition, we also generated lines that combined either 2 different gRNAs using double dU6 promoters or 4 gRNAs expressed as a single transcriptional tRNA-gRNA array (Port et al. 2014, Port and Bullock 2016). The *βtub85Dtub85D-cas9* and each gRNA line were crossed to obtain trans-heterozygous males expressing both Cas9 and the gRNA in the germline where activity is expected to occur. Next, these individuals were crossed to wild type females to determine the sex-ratio of their offspring (Figure 1C). As a control, we either used *βtub85Dtub85D-cas9* alone or the reciprocal cross that, due to the lack of *βtub85Dtub85D* activity in females, was not expected to express Cas9. Figure 1D summarizes the results of these experiments performed with a pool of individuals or by crossing single males to three females. We noted that while most gRNAs targeting X-linked repeats did not - substantially affect the sex-ratio, the *Muc14a_6* gRNA induced a highly significant male-biased sex-ratio of 61.5% (p < 0.0001). This level of distortion was consistently observed in the follow-up experiments. Of those gRNAs targeting putative X-linked haplolethal genes, two gRNAs, *RpS6_2* and *Rps5a_1*, yielded a significant excess of male progeny with frequencies of 93.8% (p < 0.0001) and 56.6% (p = 0.0112), respectively. All genetic cross data are provided in Supplementary Dataset 1.

We first focused our attention on the *Muc14a_6* gRNA that we hypothesized to induce sex-ratio distortion by the X-shredding mechanism. The *Muc14a_6* gRNA maps to the *Mucin 14A* (*Muc14a*, CG32580), an accessory-gland gene of unknown function that spans 52.6 kilobases and contains extended regions of tandemly repeated sequences. The target repeats fall within the coding sequence with a predicted 271 nucleotides between individual *Muc14a_6* gRNA target sites. We tested four gRNAs, each targeting different repeat sequences in the *Muc14a* gene and found that only one yielded significant sex distortion. *Muc14a_6* gRNA produced an excess of male progeny even if its target was not predicted to be the one with the highest number of hits in the cluster (Figure 1B). To exclude the possibility that a loss of *Muc14a* gene function is the cause of the sex-ratio distortion, we also designed two gRNAs, *Muc14a_1* and *Muc14a_2*, targeting putative non-repetitive regions in the *Muc14a* coding sequence. These experiments showed that the function of the *Muc14a* gene was not responsible for the male-biased sex-ratio observed in the progeny (Figure 1D). We performed *in-vitro* Cas9 cleavage assays and our results suggested that all gRNAs targeting repeat sequences within the Muc14a locus were active *in-vitro* and that a typical repeat unit appears targetable by each of the four *Muc14a* gRNAs (Figure 2A) more than once. We therefore concluded that the observed difference in their effects must be due to (i) different *in-vivo* performance, (ii) the particular nature or context of the target sequence or (iii) the specific repair outcomes triggered. We next tested combinations of gRNAs targeting the same or different X-linked repeat clusters to evaluate whether these could improve sex distortion rates (Figure 2B). While we observed a modest boost by providing an additional Cas9 source, neither combination of gRNAs including the transcriptional array of four gRNAs resulted in a higher distortion than that of the single *Muc14a_6* gRNA. We also observed that the *w_ex3_2* gRNA targeting *white*, when encoded as the ultimate of four gRNAs within the array, showed a dramatically reduced level of activity (Figure 2B). In flies expressing both *Muc14a_6* and *hydra_1* gRNAs from separate loci we observed comparable distortion to that of *Muc14a_6* gRNA alone. Interestingly, we observed no sex-ratio distortion when *nos*-*cas9* males, expressing the *Muc14_6* gRNA, were crossed to wild type females. This result is in stark contrast with *nos-cas9* showing substantially higher levels of activity than *βtub85Dtub85D-cas9* when combined with the *white* gRNA (Figure S1). As previously hypothesized (Windbichler, Papathanos, and Crisanti 2008, Galizi et al. 2014), these findings support the idea that X-shredding is strictly dependent on the meiotic activity of the endonuclease. An analysis of *Drosophila* development in the progenies of *βtub85Dtub85D-cas9/+, βtub85Dtub85D-cas9/Muc14a_6* and wild type fathers confirmed that the *Muc14a_6* gRNA induced no significant zygotic lethality - over any fitness effect of *βtub85Dtub85D-cas9* alone - an outcome expected if the loss of females is due to X-shredding acting pre-zygotically (Figure 2C).

**Figure 2.**
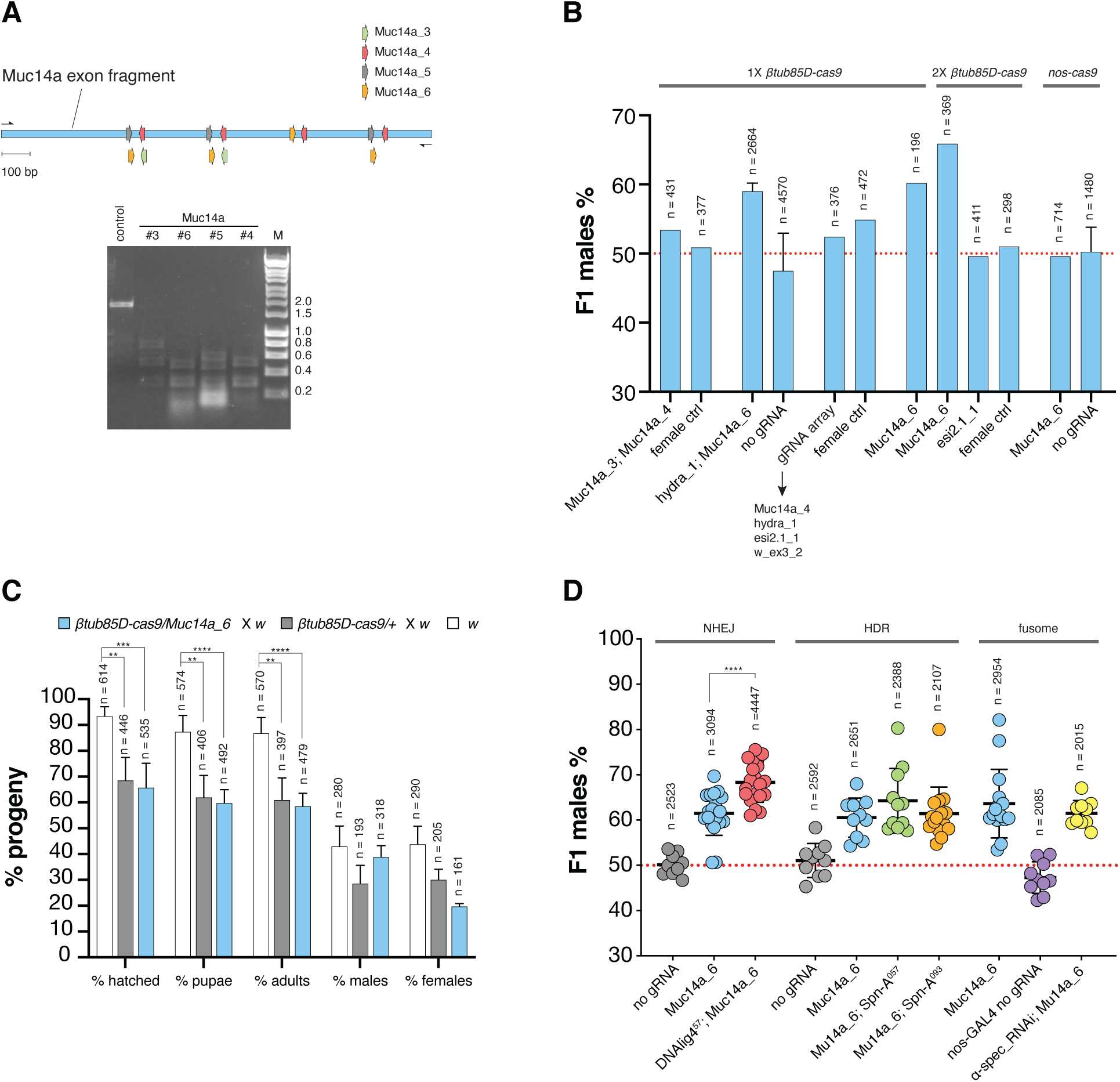
**(A)** Assay to detect CRISPR/Cas9-mediated cleavage *in vitro*. A typical region of the Muc14a gene containing at least 2 binding sites for each of the gRNAs: *Muc14a*_3, *Muc14a_4*, *Muc14a_5* and *Muc14a_6* (top). The PCR amplified DNA fragment was used as a digestion target for Cas9/gRNA cleavage reactions *in vitro* (bottom). Reactions were run on a gel to detect cleavage. A control without gRNA was included. **(B)** Analysis of combinations of gRNAs and Cas9 sources for X-shredding. Average male frequencies in the F1 progeny are shown for each parental genotype with a single copy of *βtub85Dtub85D-cas9* transgene (1X), two copies of *βtub85Dtub85D-cas9* transgene (2X) or one copy of *nos-cas9* (grey bars). All lines were crossed to wild type *w* individuals. The reciprocal cross (female ctrl) or heterozygote *βtub85Dtub85D-cas9/*+ or *nos-cas9/*+ without gRNA (no gRNA) were used as control. The black arrow indicates gRNAs in the multiplex array and the red dotted line indicates an unbiased sex-ratio. Crosses were set as pools of males and females or as multiple male single crosses in which case error bars indicate the mean ± SD for a minimum of ten independent single crosses. For all crosses n indicates the total number of individuals (males + females) in the F1 progeny counted. **(C)** Developmental survival analysis of the F1 progeny of *Muc14a_6/βtub85Dtub85D-cas9* males crossed to *w* females compared to *w* and *βtub85Dtub85D-cas9/*+ control males crossed to *w* females. n indicates the number of individuals recorded at every developmental stage (males + females) in the F1 progeny. Bars indicate means ± SD for at least ten independent single crosses. Statistical significance was calculated with a *t* test assuming unequal variance. ^**^p < 0.01, ^***^p< 0.001 and ^****^p<0.0001. **(D)** Influence of DNA repair and fusome integrity on X-shredding. Male frequencies in the progeny of *DNAlig4*^*57*^ (red), *spn-A*^*057*^ (green) and *spn-A*^*093*^ (orange) mutants and *UAS_α-spectrinRNAi*; *nos-GAL4* (yellow) knockdown mutants. The F1 seX-ratio of the progeny of males carrying *Muc14a_6/βtub85Dtub85D-cas9* in the mutant backgrounds was compared with those of *βtub85Dtub85D-cas9/*+ (grey)*, Muc14a_6/ βtub85Dtub85D-cas9* (cyan) and *nos-GAL4/*+ (purple) control males. Each dot represents the percentage of F1 males from a cross between one male and three females. n is the number of individuals (males + females) in the F1 progeny. Bars show means ± SD for at least ten independent single crosses. Statistical significance was calculated with a *t* test assuming unequal variance. ^****^p<0.0001.

To better understand the cellular mechanisms that influence the outcome of X-shredding, i.e. loss of X-bearing gametes and a bias towards males in the progeny, we generated lines bearing mutations in DNA repair pathway components or core components of the fusome. The *DNA Ligase IV* gene (*lig 4*; CG12176) encodes an ATP-dependent DNA ligase involved in non-homologous end joining (NHEJ) DNA repair; *spindle-A* (*spn-A*; CG7948) is a *Drosophila* homolog of the Rad51 gene required for double-strand break (DSB) repair by homologous recombination (HR) in both somatic and germ cells and α-*spectrin* (α-*spec*; CG1977) encodes for a component of the fusome, an organelle that facilitates intracyst cell communications during spermatogenesis (Romeijn et al. 2005, Staeva-Vieira, Yoo, and Lehmann 2003, Lu and Yamashita 2017). The fusome has also been implicated in the coordinated intracyst cell death response following DNA damage and mediating protein transport and diffusion between connected sperm cells, a process on which X-shredding in hemizygote individuals may rely on (Lu and Yamashita 2017, Windbichler, Papathanos, and Crisanti 2008). We found that the disruption of NHEJ repair by the mutant *lig4*^*57*^ significantly increased the level of male-bias to above 68%, whereas *spn-A* mutants and α-*spec* dsRNAi did not affect the sex-ratio significantly (Figure 2D).

To gain a more accurate understanding of the DNA repair mechanisms acting during X-shredding we performed sequencing of the *Muc14a* repeat cluster before and after it had undergone modification by Cas9. We crossed the *Muc14a_6* gRNA to both, *βtub85Dtub85D-cas9* and *nos-cas9* transgenes to examine the outcomes of activity at different stages of spermatogenesis. However, since F1 females inherit modified X-chromosomes from their fathers and unmodified X-chromosomes from their mothers, it is difficult to assess the exact level of gene editing on paternal X-chromosomes. To overcome this obstacle, we crossed *βtub85Dtub85D-cas9/Muc14a_6* or *nos-cas9/Muc14a_6* male individuals to X^X/Y females with attached-X-chromosomes (Kaufman 2017). By doing this, single modified X-chromosomes are passed from fathers to sons and can be analysed by amplicon sequencing (Figure 3A). In control males, ~90% of repeats contained the full gRNA target site (Figure 3B, panel 1) although polymorphisms in the surrounding sequence indicated that the Muc14a cluster shows substantial heterogeneity. We found that, on average, X-chromosomes inherited from *βtub85Dtub85D-cas9/Muc14a_6* fathers showed a > 40% reduction of cleavable gRNA target sites (Figure 3B, panel 1) but surprisingly this value was over 85%, on average, in those X-chromosomes derived from *nos-cas9/Muc14a_6* males, despite the fact that no sex-ratio distortion had been observed with this combination. X-chromosomes of *βtub85Dtub85D-cas9/Muc14a_6* fathers had, on average a larger number of unique alleles (identified from mapping reads that lacked the gRNA at the target site) than the wild type. By contrast, the progeny of *nos-cas9/Muc14a_6* showed a reduction of allele diversity, when considering alleles represented in ≥ 1% of mapping reads (Figure 3B, panel 2). A similar pattern was observed when we considered alleles with a read coverage above 10X, where X-chromosomes derived from *βtub85Dtub85D-cas9/Muc14a_6* males showed a more diverse repeat landscape compared to control and *nos-cas9/Muc14a_6* males (Figure 3B, panel 3). We analysed in more detail the fate of pre-existing alleles in the control males (Figure 3C) as well as novel alleles presumably generated by CRISPR/Cas9 activity (Figure 3D). While consensus allele frequency decreased in males of the experimental groups, we found a pre-existing allele predicted to be cleavage-resistant (AACAaATCAAGAGGAAACATCaG, with mutations in both the target site and the PAM), to increase in median frequency from <1% to ~25% in males from *βtub85Dtub85D-cas9/Muc14a_6* but not *nos-cas9/Muc14a_6* fathers (Figure 3C, row 3). In contrast, novel CRISPR-induced alleles, likely generated pseudo-randomly, were commonly shared amongst a few males, although their mapping reads accounted for more than 40% of all reads in some males from *nos-cas9/Muc14a_6* fathers (Figure 3D). On the sequence level, CRISPR-induced alleles consisted mainly of smaller deletions at or around the site of cleavage and all would be predicted to prevent further Cas9 cleavage. Together these findings suggest quite different dynamics of repair or repair mechanisms between the early acting *nos-cas9* and the meiotic *βtub85Dtub85D-cas9*.

**Figure 3.**
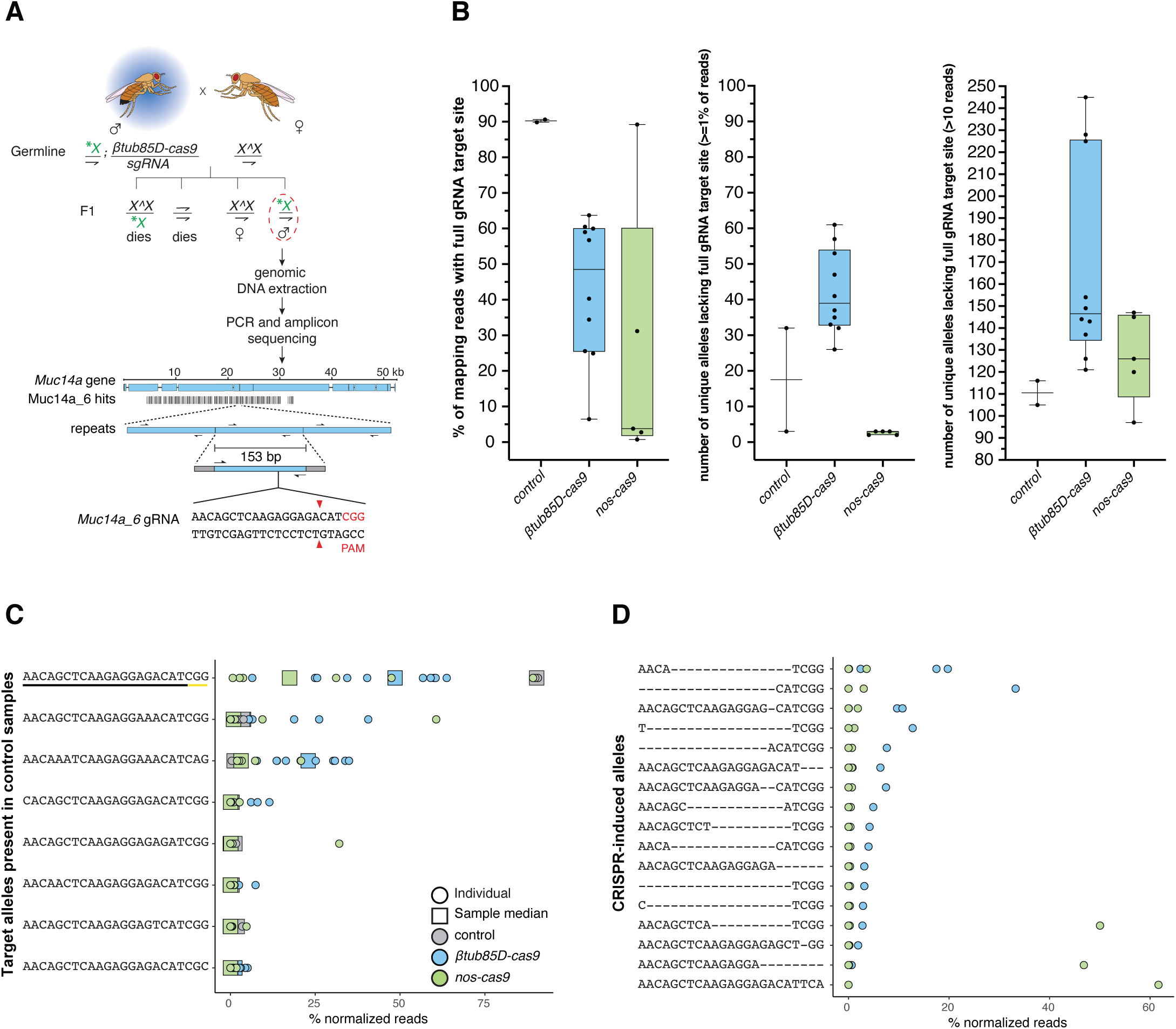
**(A)** Schematic of the genetic crosses to obtain shredded X-chromosomes from *Muc14a_6/βtub85Dtub85D-cas9* or *Muc14a_6/nos-cas9* fathers for amplicon sequencing. Trans-heterozygous males for *cas9* and the *gRNA* were crossed to females with attached-X-chromosomes (X^X). Offspring with supernumerary or lacking X-chromosomes are inviable leaving X^X/Y females and males carrying the X-shredded chromosome (patroclinous inheritance) which we selected and analysed. Genomic DNA from single males was extracted and a 153 bp DNA motif containing the *Muc14a_6* gRNA target site was amplified with primers containing Illumina Sequence adapters. As a comparison we used DNA from *βtub85Dtub85D-cas9*/+ individuals lacking the gRNA. The shredded X-chromosome is indicated by a green asterisk. A dashed red circle indicates males selected for amplicon sequencing. Vertical black bars represent the number and location of the *Muc14a_6* gRNA repeats within the *Muc14a* gene. **(B)** Analysis of allele variation at the *Muc14a_6* target site by amplicon sequencing. Indicated is the percentage of all mapping reads that contain the complete, unaltered *Muc14a_6* target site (left panel) in the control and experimental males. The middle and right panel show the number of reads harbouring all other unique alleles that represented at least 1% of all reads or were represented by 10 or more reads, respectively. **(C)** Analysis of unique alleles (including the wild type target site) at the *Muc14a_6* target site that pre-existed in both control samples. Indicated are the relative frequencies of these alleles in each control and experimental male (circles) including the median frequency of each allele in all *Muc14a_6/βtub85Dtub85D-cas9* or *Muc14a_6/nos-cas9* males (squares). **(D)** Analysis of *de-novo* alleles at the *Muc14a_6* target site not present in control males and are putative CRISPR-induced alleles. In C and D we considered here only alleles that represented ≥1% of normalized reads in at least one male sample.

We next focussed our analysis on the set of gRNAs targeting putative haploinsufficient single-copy genes. As we observed for gRNAs targeting the Muc14a repeats, no combination of gRNAs with *βtub85Dtub85D-cas9* was found to increase the sex-ratio relative to the *RpS6_2* gRNA alone. Both gRNAs, *RpS5a_1* and *RpS6_2*, when crossed to *βtub85Dtub85D-cas9* were individually capable of inducing male-bias sex-ratio distortion, but when combined within a gRNA array, we found a significantly lower level of distortion in the progeny than when we used *RpS6_2* alone. In contrast to X-shredding, X-meddling is expected to act post-zygotically by inducing female lethality in the developing embryo likely as a result of an insufficient dose of the target-gene product. We examined *Drosophila* development in the progenies of *βtub85Dtub85D-cas9/*+, *βtub85Dtub85D-cas9/RpS6_2* and wild type fathers and, in contrast to the *Muc14a_6* gRNA, we did observe significant lethality at the embryonic and subsequent stages. This result would suggest a mechanism of lethality that operates during development and that the few survivor females in the progeny carried rescue mutations that restored the function of the ribosomal target genes, while preventing further CRISPR cleavage. To confirm this hypothesis, we performed amplicon sequencing of the *RpS6_2* target sequence using 2 pools of the surviving females from *βtub85Dtub85D-cas9* fathers expressing the *RpS6_2* or *RpS6_2* and *RpS6_1* gRNAs (Figure 4C). Surprisingly, while we did observe possible rescue alleles, the majority of mutant alleles identified in surviving females are not expected to restore *RpS6* gene function because of the presence of translational frameshifts (Figure 4D). Although, one has to be cautious to infer loss of function from such experiments alone (Smits, A.H et al. 2019), these results imply that the mechanism of lethality following *RpS6* cleavage and imperfect repair is dominant rather than dependent on an insufficient gene dose. Finally, a combination of *Muc14_6* and *Rps6_2* gRNAs, i.e. combining the two mechanism of distortion we have described, yielded the highest level of distortion we observed in this study (on average 95.8% males) although this was not significantly different from *RpS6_2* alone.

**Figure 4.**
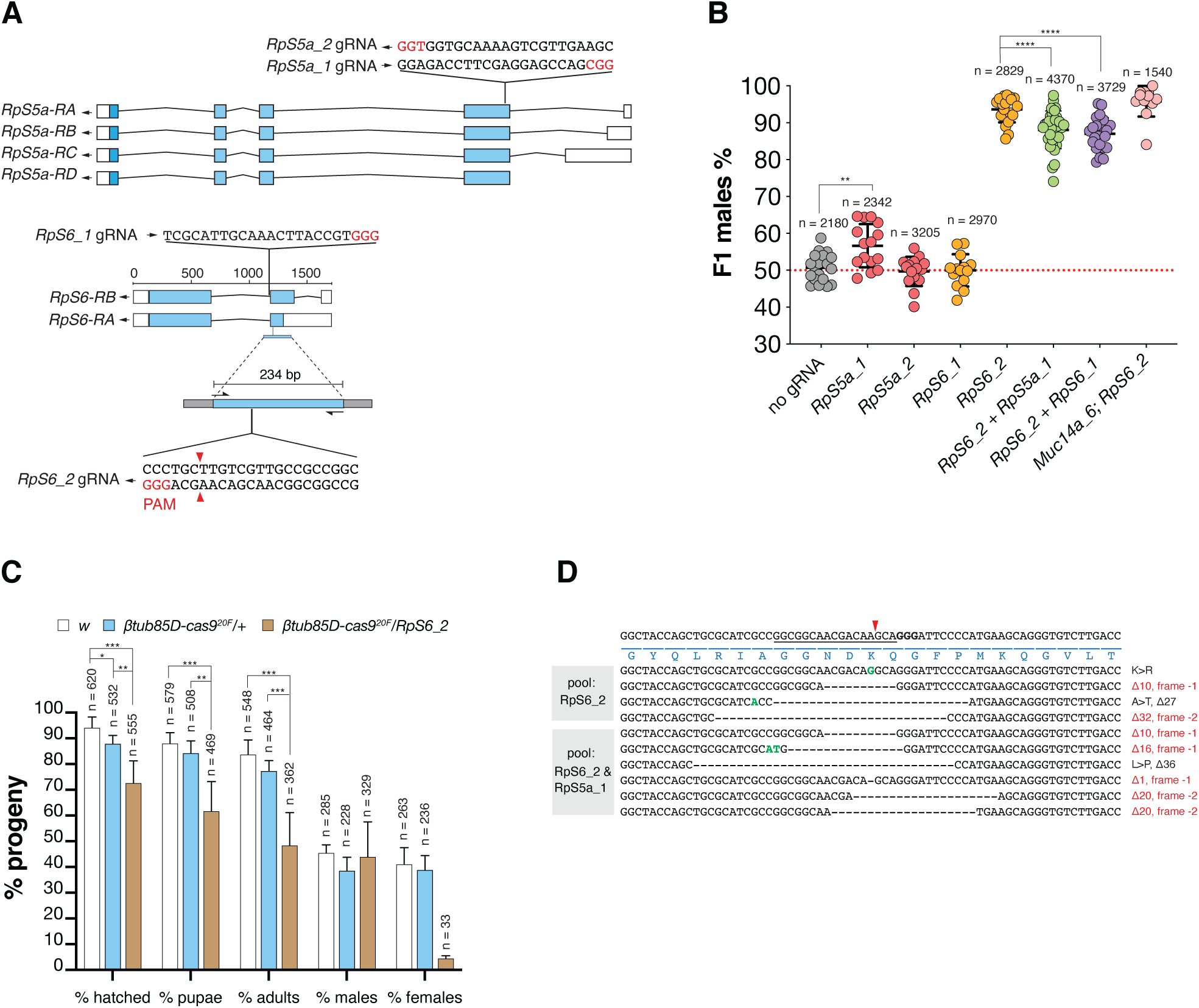
**(A)** gRNA target sites within presumptive haploinsufficient genes on the X-chromosome. Schematic representation of *RpS5a* and *RpS6* gene organization. Both genes encode for a small ribosomal subunit protein (RpS). As illustrated in the figure, the *RpS5a* gRNAs, *RpS5a_1* and *_2*, map in the fourth exon shared by all four transcripts of the gene and the *RpS6* gRNAs*, RpS6_1* and *2*, map in the third exon of two transcripts in the corresponding gene. The figure shows the 234 bp fragment (with Illumina Sequence adapters on both sides as grey boxes) surrounding the *RpS6*_2 gRNA target site that was used for amplicon sequencing. Blue boxes indicate coding sequences, white boxes indicate UTR regions and PAMs are indicated in red. **(B)** Efficiency of single gRNAs or combinations of gRNAs for X-meddling. Shown is the frequency of males in the progeny from *βtub85Dtub85D-cas9* males combined with four *gRNA* lines and crossed to wild type *w* females. Individual gRNAs, *RpS5a_1* or *RpS5a _2 (*red), and *RpS6_1* or *RpS6_2* (orange) are compared to double gRNA arrays co-expressing *RpS6_2* + *RpS5a_1* (green) or *RpS6_2* + *RpS6_1* (purple) as well as a combination of both *RpS6_2* and *Muc14a_6* transgenes (pink). As a control, crosses from *βtub85Dtub85D-cas9/*+ (no gRNA, grey) fathers are shown. Each dot represents the percentage of F1 males from a cross between one male and three females. n is the number of individuals (males + females) in the F1 progeny. Black bars show means ± SD for at least ten independent single crosses. Statistical significance was calculated with a *t* test assuming unequal variance. ^**^p < 0.01, ^***^p <0.001 and ^****^p<0.0001. **(C)** Developmental survival analysis of the F1 progeny of *RpS6_2/βtub85Dtub85D-cas9*, *w* or *βtub85Dtub85D-cas9/*+ males crossed to *w* females. n is the number of individuals recorded at every developmental stage (males + females) in the F1 progeny. Bars indicate means ± SD for at least ten independent single crosses. Statistical significance was calculated with a *t* test assuming unequal variance. ^**^p < 0.01, ^***^p<0.001 and ^****^p<0.0001. **(D)** CRISPR induced target site mutations within the *RpS6* gene analysed by pooled amplicon sequencing of surviving female progeny. On top, the wild type DNA sequence spanning the *RpS6_2* gRNA target site in the *RpS6* ribosomal gene is shown with the gRNA binding position (underline), the cut site (red arrowhead), the PAM (bold nucleotides) as well as the encoded amino acids (blue). Target site variants identified in pools of F1 females from *RpS6_2/βtub85Dtub85D-cas9* fathers or *RpS6_2 RpS5a_1/βtub85Dtub85D-cas9* fathers are shown below. Dashed lines correspond to nucleotide deletions, green coloured bases represent insertions. Predicted amino-acid substitutions (>) are indicated to the right.

## Discussion

To recreate in *Drosophila* a synthetic X-shredding mechanism pioneered in *A. gambiae*, we generated gRNAs targeting repeat sequences we identified on the X-chromosome. This demonstrates that X-shredding is a transferable mechanism and not linked to the particular nature of the mosquito target, i.e. the ribosomal DNA gene cluster that was targeted in previous studies (Galizi et al. 2014, Galizi et al. 2016). It also shows that CRISPR/Cas9 target sequences that are able to induce sex-ratio distortion can be identified bioinformatically. To search for such X-linked repeats, we employed the Redkmer pipeline using raw sequence data as the only input. This is crucial as many target species of medical or agricultural importance are likely to lack high-quality genome assemblies. Even when such assemblies exist, repeats sequences, a problematic class of sequences for assemblers, often remain poorly resolved. However, recent progress in telomere-to-telomere chromosome assemblies that incorporate large DNA repeat clusters may simplify this step even further in the future (Dilthey et al. 2019). The level of sex distortion towards males we observed in the fly was not as extreme as observed in *A. gambiae*. Given that activity against the single-copy *white* was incomplete this suggests possible improvements by further optimizing the level of Cas9 expression in the germline of *Drosophila*. Using our best Cas9 strain, we found that achieving higher rates of distortion required us to interfere with mechanisms of DNA repair. This in turn indicates that targeting more repetitive sequences could be another avenue to enable a more dramatic bias towards males, though the *Drosophila* X-chromosome lacks such repeats. The sequence microenvironment rather than the number of repeats may also play a role in determining why certain gRNAs trigger gamete loss while others don’t. The identification of X-chromosomes which are more suitable for X-shredding in target species of medical or agricultural relevance is thus the next task. Our study provides clear lessons for the application of X-shredding to such species. A strong meiotic promoter yielding high levels of Cas9 should be combined with as few gRNAs as possible, ideally one, targeting a highly repetitive sequence, which does not have to be essential and can consist of a single repeat cluster on the X-chromosome.

Chromosome-wide, distributed repeats represent an alternative set of targets but the lack of such sequences in *Drosophila* precluded us from evaluating their use for X-shredding. The fly literature suggests that the 1.688 X-chromosome satellite involved in X dosage compensation (Menon et al. 2014) one of the most abundant repetitive sequences in *Drosophila melanogaster*, would represent such a target (Kim et al. 2018). While the candidate did include a number of kmers that could be attributed to the 1.688 satellite they did not pass our selection criteria or were significantly less abundant than the targets we selected. This is likely due to the heterogeneity and stratification of 1.688 repeats in various chromosomal locations (Kuhn et al. 2012). Although the use of larger arrays of gRNAs has been suggested to compensate for the lack of abundant X-linked repeats (Galizi et al. 2016), our data suggests that targeting a highly repetitive sequence with a single gRNA may be a more viable route. Even if multiple gRNAs could be expressed efficiently and concomitantly, they could compete for access to Cas9 protein in the loading step thus reducing activity of each individual gRNA.

Our data suggests that the high level of activity of *nos-cas9* with *Muc14a_6* gRNA during early spermatogenesis could cause the repeats to be subject to multiple cleavage-repair cycles which in turn may lead to the observed reduction in the complexity and presumably also the size of the repeat cluster (and no sex bias). By contrast, *βtub85Dtub85D-cas9/Muc14a_*6 activity, which acts later in meiosis, would encounter, following multiple mitotic divisions, more X-chromosomes to target in the larger population of primary spermatocytes on which it may be acting continuously and in parallel to generate novel alleles. This would result in a broader spectrum of mutations inherited by the progeny. Alternatively, the observed differences could also partly relate to the predominance of homologous over non-homologous DNA repair pathways acting with varying stringency during the early (including stem cells) and late stages of spermatogenesis, respectively (Chan et al. 2011). While *nos-cas9* would favour the loss of repeat units by recombination-based repair mechanism, *βtub85Dtub85D-cas9* would trigger NHEJ repair events leading to a more diverse allele pool. Indeed, we found little evidence of homologous repeat to repeat repair in *βtub85Dtub85D-cas9/Muc14a_6* males. For instance, the pre-existing cleavage-resistant allele AACAaATCAAGAGGAAACATCaG is associated with a 6bp indel polymorphism located 53 nucleotides upstream of the gRNA target. We did not detect dissociation of this SNP even in samples in which the cleavage resistant allele rose to a frequency of 30% of the read pool. Dissociation would have indicated that a double-strand break in a wild-type repeat unit had been repaired using the cleavage resistant allele as a template.

One caveat is the fact that the X-chromosomes from *βtub85Dtub85D-cas9/Muc14a_6* males we analysed by amplicon sequencing managed to escape the germline in the form of X-bearing gametes and gave rise to viable males. One might argue that they represent chromosomes with an overall lower level of modifications or modifications with repair outcomes compatible with transmission and male survival (which may in turn differ from the requirements for female survival). In contrast, the X-chromosomes unable to form viable gametes, i.e. those that we could not analyse by sequencing the surviving progeny, may be the ones that underpin the sex distortion observed, and may be subject to fundamentally different repair events.

A number of questions remain to be answered, such as the mechanism through which X-shredding causes the loss of X-bearing gametes and how insufficient or incomplete DNA repair is associated with this process. This outcome is by no means self-evident; for example, it has been shown that functional sperm can be produced despite lacking either one or both major autosomes or lacking DNA (Lindsley and Grell 1969). The fly model of X-shredding we have established will allow to tackle these questions experimentally.

We also established an X-meddling system in the *Drosophila* model by targeting X-linked ribosomal genes and achieved high rates of male bias. This latter strategy, in the form of a Y-linked endonuclease, has recently been proposed as an efficient alternative to gene drives for genetic control (Burt and Deredec 2018). Modelling suggests Y-linked editors would outperform other self-limiting strategies while having less impact on non-target populations when compared to gene drives or driving Y-chromosomes. We found the mechanism of embryo lethality in the case of the ribosomal *RpS6* gene to be more complex than anticipated. Rather than protein dose insufficiency, a dominant lethal effect may partially or totally explain our results. Such an effect in mutated ribosomal proteins has been observed previously (Devlin et al. 2010) and could, for example, be explained by the poisoning of ribosomes with dysfunctional ribosomal proteins. Since, in this case, every escaper female represents a single repair event, phenotypes would be expected to vary depending on the outcome of DNA repair. Further modelling may be required to understand how these phenotypes would impact genetic control at the population level. The X-shredding and the X-meddling transgenes we have described here could be expressed from the *Drosophila* Y chromosome which has recently been modified to harbour pre-characterized gRNA sites for transgene insertion (Buchman and Akbari 2019). This could allow, for the first time, to explore the use of such Y-linked editors with gRNAs for X-meddling or X-shredding. To what degree these two strategies could be designed around or would be susceptible to X-chromosome inactivation in the male germline of *Drosophila* is a further research question, in particular as this process is not fully understood in the fly (Vibranovski et al. 2012, Mikhaylova and Nurminsky 2011).

The application of X-meddling to agriculturally or medically relevant species would appear to be a more straightforward proposition in particular as ribosomal target genes show high levels of conservation. Also, the characterization of Y chromosomes and the identification of male determining regions (Meccariello et al. 2019, Krzywinska et al. 2016, Hall et al. 2015) and docking sites (Buchman and Akbari 2019) within which to land transgenes has recently made great progress.

## Acknowledgements

This study was funded by the BBSRC under the research grant BB/P000843/1 to N.W.

P.A.P. was funded by the Italian Ministry Education, University and Research (MIUR—D.M. no. 79 04.02.2014), by the United States – Israel Binational Agricultural Research and Development Fund (Research Grant No. IS-5180-19) and by the Israel Science Foundation (Research Grant No. 2388/19). The authors would like to thank Steve Russell and Adam Phillippy for helpful discussions and for providing error-corrected whole genome sequence data. We would like to thank Alex Nash, Paolo Capriotti and Rita Colonna.

## Supplementary Figure Legends

**Figure S1.**
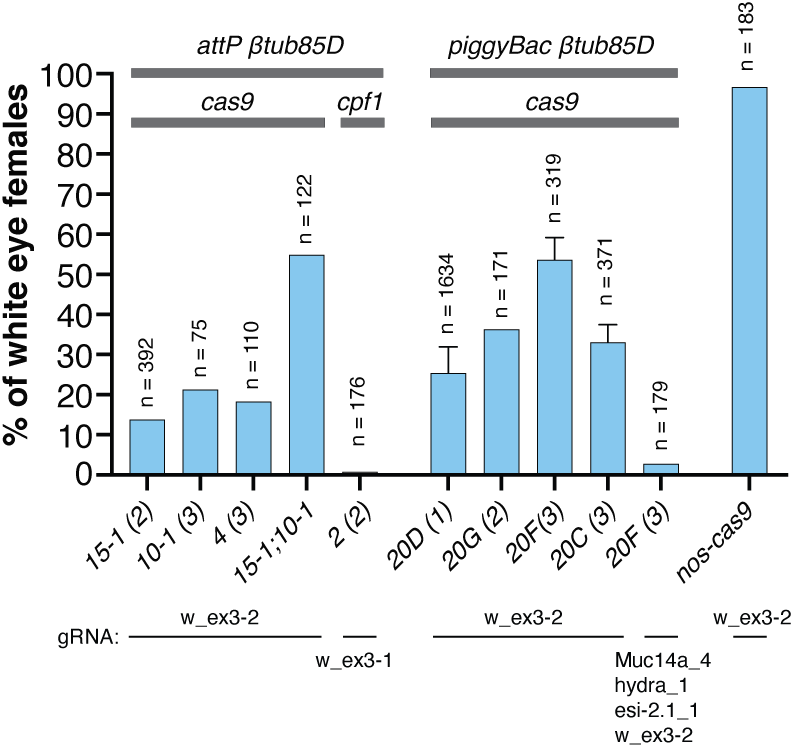
Evaluation of Cas9 activity when expressed from *βtub85Dtub85D-cas9* and *βtub85Dtub85D_cpf1* transgenes integrated at various genomic locations using ϕC31 (C31 (*attP)* or *piggyBac* mediated transformation. Flies carrying *βtub85Dtub85D-cas9* expressed from *attP* docking #15-1 on the second and #10-1 and #4 on the third chromosomes and from piggyBac mediated random integrations #20D on the X, #20 G on the second and #20F and #20G on the third chromosomes were crossed to lines transgenic for *w_ex3-2* gRNA that targets the *white* gene on the X-chromosome. *βtub85Dtub85D-cas9*/*w_ex3-2* F1 males with red eyes were then crossed to *white* mutant females, and the female progeny scored for white eyes. Experiments combining two *βtub85Dtub85D-cas9* transgenes with the *w_ex3-2* gRNA and *βtub85Dtub85D-cas9* with *w_ex3-2* expressed as part of a gRNA multiplex array were also performed. Similarly, *βtub85Dtub85D_cpf1/w_ex3-1* red eye males were crossed to *white* females, and the female progeny scored for white eyes. Along with the *βtub85Dtub85D* promoter, the *nanos*-*cas9* transgene efficiency was tested with the same *w_ex3-2* gRNA. gRNAs used for each experiment are shown below the graph. n is the number of individuals (males + females) in the F1 progeny.

**Figure S2.**
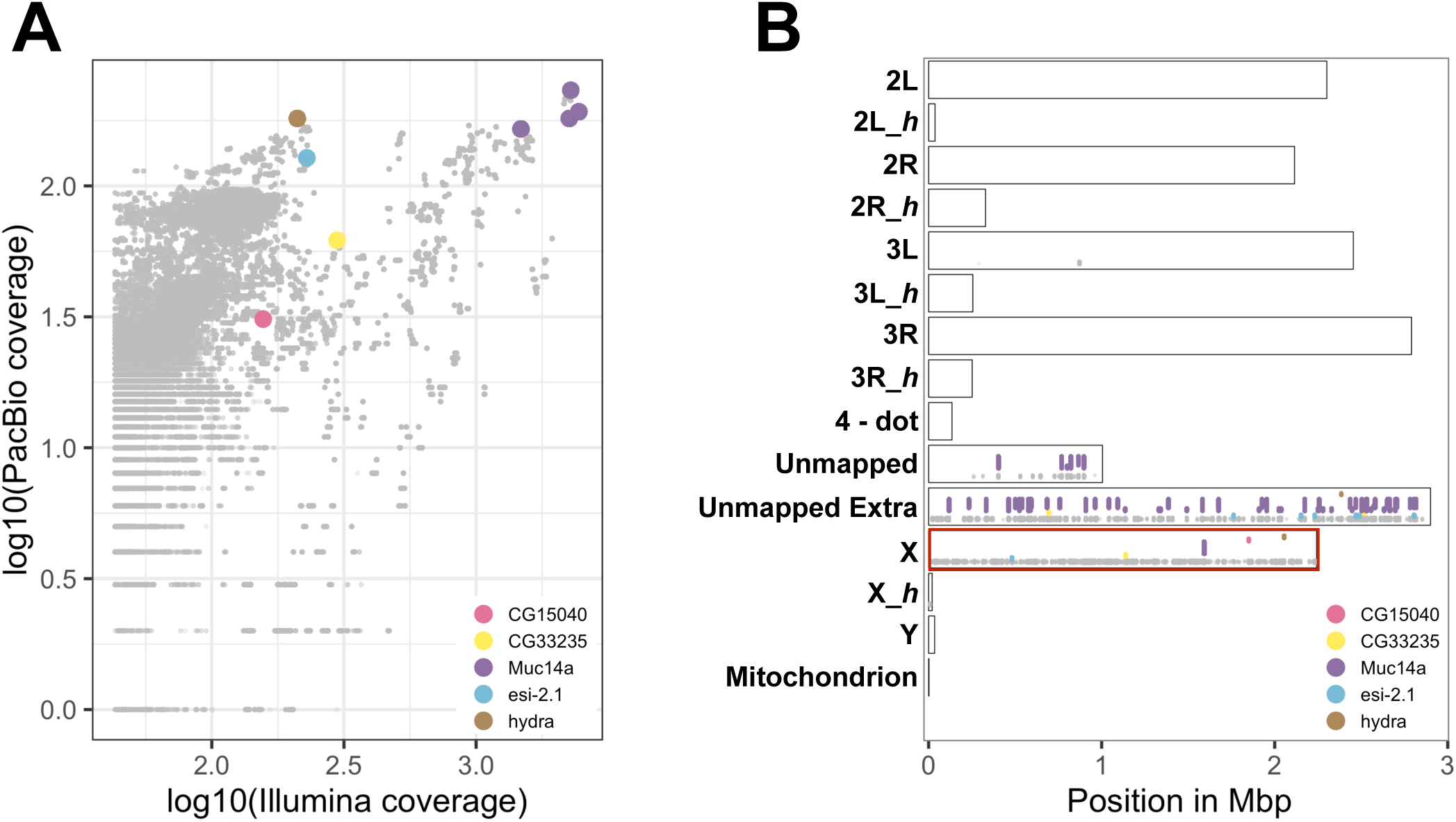
Candidate X-kmer abundance and chromosomal distribution. **(A)** Coverage of all candidate X-kmers (grey dots) including those X-kmers chosen for experimental evaluation (colored dots) within the Illumina and PacBio whole-genome sequencing read datasets as predicted by Redkmer. **(B)** Genomic distribution of all candidate X-kmers (grey dots) and those selected X-kmers for experimental testing (colored dots) on *D. melanogaster* chromosome arms based on perfect complementarity. The X-axis indicates the Mbp position of matches on each chromosomal arm, including the 4^th^ chromosome, unmapped contigs and heterochromatin (_*h*).

**Figure S3.**
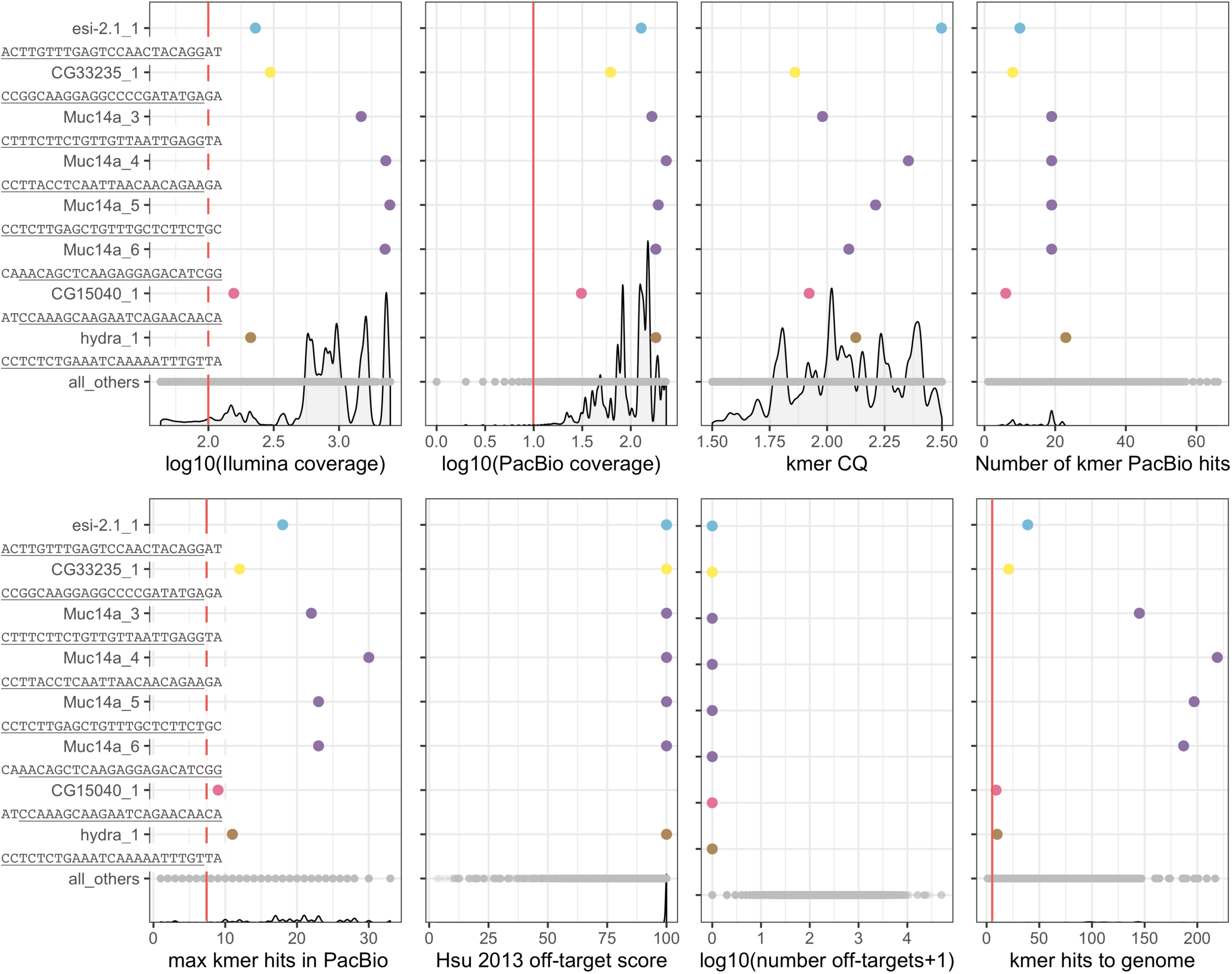
Kmer sub-selection applied to Redkmer output. Criteria for candidate kmer selection (X-axes) are shown for each of the eight selected X-kmers for X-shredding (colored dots) and for the remaining candidate X-kmers (grey dots). Red vertical lines highlight the minimum cutoff values imposed for the final target site selection. Density plots of each criteria are also shown for the entire Redkmer candidate X-kmer output. The part of the kmer sequence that represents the target sites of experimental gRNAs is indicated (underline).

## Materials and Methods

### Identification of candidate kmers for X-shredding

Error-corrected PacBio reads derived from males of the ISO1 strain (ref doi: 10.1038/nbt.3238) were obtained from the University of Maryland Center for Bioinformatics and Computational Biology (http://gembox.cbcb.umd.edu/mhap/data/dmel.polished.fastq.gz). Illumina reads from males and females of the same strain were obtained from the NCBI short read archive (female: SRX826515 and male: SRX826516). These were used as inputs to run Redkmer (ref) The mitochondrial genome (NCBI accession number: NC_024511) was used to filter out mitochondrial derived reads. We only considered Pacbio reads between 2kb and 100kb to improve chromosome quotient accuracy (CQ - Hall et al 2016) and excluded kmers occurring less than 4 times in the combined male and female Illumina data. We applied a number of critera to the 25,298 candidate X-kmers predicted by redkmer to be X-linked abundant sequences. First, we used FlashFry (McKenna et al 2018 10.1186/s12915-018-0545-0) to select among the candidate X-kmers, those that contained a sequence suitable for the design of a Cas9 gRNA. To estimate off-target potential, we used as a reference all PacBio reads predicted by Redkmer to represent autosomal and Y derived sequences. We then applied several additional metrics such as the maximum number of occurrences of a kmer per PacBio read, giving an estimate of the repeat size, and the total number PacBio reads containing the kmer at least once, giving an estimate of the abundance of the repeat across the X-chromosome. Kmers shortlisted for testing were selected based on the following cutoffs: abundance in the genome (coverage in combined male and female Illumina data >log_10_(2), coverage in PacBio data >log_10_(1), the maximum kmer hits per PacBio read > 7.5, and at least 5 perfect blast hits to the assembled genome. All candidate X-kmers were also re-assembled into longer contigs using the Geneious Software (Geneious Assembler allowing no mismatches) to identify possible higher-order repeat loci and to ensure that different sequence classes would be targeted in our experiments. A total of 8 of 205 X-kmers passing all filters were then selected for testing in transgenic strains.

### Generation of transgenic lines

#### Design and assembly of constructs

Unless otherwise noted, cloning was performed with the NEBuilder Hi Fi DNA Assembly kit (New England Biolabs). PCR reactions were performed with the Phusion High-Fidelity PCR Master Mix with HF buffer (New England Biolabs). All inserts were verified by sequencing (GENEWIZ). Primers used for plasmid construction are listed in Table1 S1.

#### *βtub85D-cas9* expression plasmid

The previously described plasmid *pYSC47w*^−^ harbouring the *3Px3-eGFP* transformation marker, *attB* and *piggyBac* recombination sequences (a gift from Andrea Crisanti, Imperial College London) was used to build *Drosophila cas9* constructs. The plasmid was linearized with *Sbf*I. A 478 bp DNA fragment of *βtub85Dtub85D* 5’ regulatory region was amplified with primers *βtub85Dtub85D F/βtub85Dtub85D-Cas9-R* from the genomic DNA of *Drosophila* strain *w*^*1118*^. The human codon-optimized Cas9 coding sequence including two nuclear localization signals (*SV40 NLS* at the 5’ and *nucleoplasmin NLS* at the 3’) was amplified from *hcas9* (a gift from George Church; Addgene plasmid # 41815; http://n2t.net/addgene:41815; RRID:Addgene_41815) using primers *βtub85Dtub85D-cas9-F/cas9-βtub85Dtub56D-3’UTR-R* (Mali et al. 2013). A third DNA fragment of 651 bp in the *βtub85Dtub56D* 3’-UTR region was amplified with primers casβtub56Ftub56F/βtub56Ftub56D R from the plasmid *pYSC61*. The three fragments were cloned in the open backbone of *Sbf*I-linearized *pYSC47w*^−^ resulting in the final plasmid termed *“pYSC47w+_βtub85Dtub85D_cas9_βtub85Dtub56D”*. The plasmid was used for both *piggyBac* and *attP*-docking site integration.

#### *βtub85D-Lbcpf1* expression plasmid

To generate the plasmid expressing *cpf1*, fragments containing the *βtub85Dtub85D* promoter and the *βtub85Dtub56D* 3’-UTR were amplified from the construct *pYSC47w+_βtub85Dtub85D_cas9_βtub85Dtub56D.* The fragment containing the *LbCpf1* coding sequence was amplified from *pY016 hLbCpf1* (Addgene plasmid # 69988; http://n2t.net/addgene:69988; RRID:Addgene_69988) {Zetsche, 2015, r01553}. The three fragments were finally assembled in *Sbf*I-digested *pYSC47w*^−^.

#### Guide RNA expression plasmid

All single guide RNAs were cloned in *Bbs*I-linearized *pCFD3-dU6:3 gRNA* plasmid (a gift from Simon Bullock; Addgene plasmid # 49410; http://n2t.net/addgene:49410; RRID:Addgene_49410) (Port et al. 2014) and harbouring a U6 promoter. A gRNA array containing *Muc14a_4*, *hydra_1*, *esi2.1_1* and *w_ex3_2*, in this order, was assembled with the oligos Array1_Cas9_PCR1_F, Array1_Cas9_PCR1_R, Array1_Cas9_PCR2_F, Array1_Cas9_PCR2_R and Array1_Cas9_PCR3_R and cloned in *pCFD5* plasmid (a gift from Simon Bullock; Addgene plasmid # 73914; http://n2t.net/addgene:73914; RRID:Addgene_73914) (Port and Bullock 2016). Two distinct arrays, each containing two gRNAs targeting ribosomal genes, were assembled with primers RpS6 *2_*pCFD4_F and RpS6_ 1_pCFD4_R for the *RpS6_2 + RpS6_1* array and RpS6_2_pCFD4_F and RpS5a_1_pCFD4_R for the *RpS6_2 + RpS5a_1* array. The arrays were cloned in *pCFD4-U6:1_U6:3* tandem gRNAs plasmid (a gift from Simon Bullock; Addgene plasmid # 49411; http://n2t.net/addgene:49411; RRID:Addgene_49411). The plasmids *pCFD3*, *pCFD4* and *pCFD5* were all integrated in *attP*-docking sites. CRISPR target site design. CHOPCHOP v2 was used to choose gRNA target sites in *RpS6* and *RpS5a* ribosomal genes specific regions in *Drosophila* genome (dm6).

#### Embryo injections

Embryo injections were carried out at the University of Cambridge Fly Injection Facility. The *βtub85Dtub85D_cas9* constructs were inserted at the *P{CaryP}attP40* site on the 2nd chromosome (25C6; Stock 13-20) and the *PBac{y+-attP-9A}VK00027* on the 3rd chromosome (89E11; Stock 13-23), both stocks marked with *yellow*^+^. Moreover, the *βtub85Dtub85D_cas9* plasmid was integrated in *D. melanogaster* by piggyBac transposition. Transgenic flies were balanced with *w*^*1118*^; *if/CyO* and *w*^*1118*^; *TM3, Sb/TM6B*. Different lines were generated with *βtub85Dtub85D_cas9* integrated on the X (20D), on the second (20G) and on the third chromosome (20F). gRNAs cloned in *pCFD3*, *pCFD4* or *pCFD5* were integrated in the genome at the *P{CaryP}attP40* site on the second (Bloomington stock 25709) and/or *P{CaryP}attP2* site on the third chromosome (Bloomington stock 25710). *pCDF4_RpS6_2_RpS6_1, pCDF4_RpS6_2_RpS5a_1, pCFD3_w_ex3-2, pCFD3_RpS6_gRNA2, pCFD3_RpS6_gRNA1, pCFD3_Muc14a_1, pCFD5_Cas9_Array1, pCFD3_CG33235_1, pCFD3_CG15040_1, pCFD3_Hydra_1, pCFD3_Muc14a _4, pCFD3_ Muc14a_3, pCFD3_ Muc14a_3, pCFD3_ Muc14a_5, pCFD3_ Muc14a_6*, and *w_ex3-1* (LbCpf1) were inserted in the *P{CaryP}attP2* site on the third chromosome and *pCFD3_RpS5a_gRNA1, pCFD3_RpS5a_gRNA2, pCFD3_ Muc14a_2, pCFD3_Hydra_1, pCFD3_ Muc14a_3, pCFD3_ Muc14a_5, and pCFD3_ Muc14a_6* were inserted in the *P{CaryP}attP40* site on the second chromosome.

### Fly husbandry and strains

Flies were maintained under standard conditions at 25°C with a 12/12 hour day and night cycle. For general maintenance, stocks were provided with new food every 2-3 weeks. Flies were anesthetized with CO_2_ during phenotyping. To assess fluorescent green eyes phenotype conferred by the *3PX3:eGFP* transgene, we used a conventional fluorescence microscope. Description of stocks not provided here can be found in FlyBase (http://flybase.org). The lines used in this study were: *DNAlig4*^*57*^ Bloomington Stock #8520, *Spn-A*^*057*^ and *Spn-A*^*093*^ a gift from Mitch McVey (Tuft University), Df(3R)XF3 Bloomington Stock #2352, *UAS-α-spectrinRNAi* Bloomington Stock #56932 (TRIP.HMC04371), *nanos_Gal4* Bloomington Stock #4937, *C(1)DX/FM6* (compound chromosome with attached X-chromosomes) Bloomington stock #784.

### Analysis of *βtub85D-cas9* and *nos-cas9* activity

To test *attP* or *piggyBac* integrated *βtub85Dtub85D-cas9* and *nos-cas9* efficiencies, females from each line were crossed to *w*^+^; *w_ex3-2* gRNA males bearing a gRNA targeting the *white* gene on the X-chromosome. Red eye trans-heterozygous *w*^+^; *cas9*/*w_ex3-2* F1 males were selected, crossed to *w*^*1118*^ females and the percentage of white eye females in the progeny was recorded.

### Genetic crosses for X-shredding and X- meddling

During the initial screen for X-shredding-induced sex ratio distortion, *βtub85Dtub85D-cas9* mothers were crossed to each gRNA line (from a 25 nucleotides kmer), in a vial at 25°C. At least twenty trans-heterozygous F 1 *gRNA/*+; *βtub85Dtub85D-cas9/*+ males for each combination of sgRNA were then crossed to the same number of *w*^−^ females in new vials at the same temperature. We used the reverse crosses, *βtub85Dtub85D-cas9/sgRNA* females crossed to *w*^−^ males, as controls due to the lack of *βtub85Dtub85D* activity in females. We discarded the gRNA lines that did not result in sex ratio distortion while those where we recorded a male bias progeny were further tested by pursuing single cross analysis. For every single cross, a single trans-heterozygous *βtub85Dtub85D-cas9/+;gRNA/*+ male (experiment) and a heterozygous *βtub85Dtub85D-cas9/*+ male (control) were crossed to three wild-type females in separate vials at 25°C. We set up a minimum of ten single crosses for each genotype analysed. For all crosses where we analysed the change in male bias in the progeny of *βtub85Dtub85D-cas9/+;Muc14a_6/*+ in a NHEJ, HDR or fusome mutant backgrounds, we set up ten-twenty replicas of single male crosses, i.e., *w DNAlig4*^*57*^; *Muc14a_6/βtub85Dtub85D-cas9*^*20F*^ single male X three *w* females, *Muc14a_6/βtub85Dtub85D-cas9*^*20G*^;*Spn-A*^*093 or 057*^/*Df(3R)XF3* single male X three *w* females and *UAS-α-spectrinRNAi/Muc14a_6; βtub85Dtub85D-cas9*^*20F*^/*nanos_Gal4* single male X three *w* females and compared the results to the progenies of *Muc14a_6/βtub85Dtub85D-cas9* and *βtub85Dtub85D-cas9/*+ single male crosses used as controls.

In all crosses for X-meddling-induced sex ratio distortion, *βtub85Dtub85D-cas9* mothers were crossed to each gRNA analysed, in a vial. Single F1 *βtub85Dtub85D-cas9/+;gRNA/*+ (experiment) and *βtub85Dtub85D-cas9/*+(control) males were then crossed to three wild-type females in separate vials. We set up a minimum of ten single crosses for each genotype to generate means and standard deviations for statistical comparisons and thus measure consistency and robustness of the results. All crosses were done at 25°C. Percent of males and females was calculated as the ratio between the number of individuals counted and the number of individuals expected for each genotype. Flies were scored and examined with the Nikon SMZ1500 stereomicroscope equipped with a CoolLED *p*E-300 Led fluorescent illumination.

### *In vitro* cleavage assay

Kmers *Muc14a_3*, *Muc14a_4*, *Muc14a_5* and *Muc14a_6* and Cas9 were tested for *in vitro* activity using the Guide-it sgRNA *In Vitro* Transcription and Screening System (Takara Bio USA, Inc.). Each kmer was transcribed and purified according to the *In Vitro* Transcription of sgRNA protocol. Two experiments were performed, allowing transcription to take place for 4 and 8 hours. Genomic DNA templates were obtained from Muc14a fragments previously cloned in a *pMiniT 2.0* vector using PCR Cloning Kit (NEB) and primers kmer Muc14_F and 58537687_F. The genomic target was amplified by PCR with Cloning Analysis Forward and Reverse Primers, mapping in the *pMiniT 2.0* plasmid. The amplicon corresponding to a genomic fragment of 1240 bp was run on a gel and the DNA was excised and purified with Monarch DNA Gel Extraction Kit (NEB). DNA concentration and absorbance ratio were measured with a NanoDrop Spectrophotometer (ThermoScientific). PCR amplification of target DNA and a Cas9 cleavage assay were then carried out according to the protocol.

### Amplicon sequencing analysis preparation

Genomic DNA was isolated from ten single F1 males originated from the cross *βtub85Dtub85D_cas9*^*20F*^/*Muc14a_6* X *C(1)DX/FM6* (compound chromosome with attached X-chromosomes) Bloomington stock #784 and from one control male from the cross *w/Y; βtub85Dtub85D_cas9*^*20F*^/ *TM6B* using the QIAamp DNA Micro Kit. Genomic loci containing the *Muc14a_6* gRNA target site were amplified with Phusion HF DNA polymerase (Thermo Scientific) using Repeat 56910823_illumina_F and Repeat_Muc14a_3_illumina_R primers containing the Illumina adapters. 200 ng of genomic DNA in a 100 μl reaction volume were used as a template for a limiting PCR reaction to amplify 153 bp of slightly different repeats in the *Muc14a* gene. To maintain the proportion of reads corresponding to particular repeats, the PCR reactions were performed under non-saturating conditions for a total of 25 cycles with 55°C annealing temperature.

For deep sequencing analysis of the ribosomal gene *RpS6* target site, the genomic DNA was extracted from a pool of four daughters from the cross *βtub85Dtub85D_cas9*^*20F*^/*Rps6_2* X *w*, four daughters from the cross *βtub85Dtub85D_cas9*^*20F*^/*Rps6_2+ RpS5a_1* X *w* and from one *βtub85Dtub85D_cas9*^*20F*^/+ daughter control. The primers RpS6_1F and RpS6_1R were used to amplify a 234 bp DNA fragment spanning the *RpS6_2* target site. The amplicons were purified with NEB Monarch PCR & DNA Cleanup Kit and quantified with Nanodrop. 200 ng were checked on a gel and 500 ng were sent to GENEWIZ to be sequenced with NGS-based amplicon sequencing. We ran CRISPResso (Pinello et al. 2016) software on raw sequencing data to detect mutations at the target site using parameter -q 30, setting the minimum average read quality score (phred33) to 30.

### Viability studies

To identify the developmental stage at which the progeny from *Muc14a_6/βtub85Dtub85D_cas9* and *RpS6_2/βtub85Dtub85D_cas9* crossed to *w*^*1118*^ die, we quantified egg hatching, pupae and adult death rates in the F1. To quantify the egg hatching rate, 20–30 heterozygous *Muc14a_6/βtub85Dtub85D_cas9* or *RpS6_2/βtub85Dtub85D_cas9* and 20-30 *w*^*1118*^ virgin females were set up in embryo collection cages with grape juice agar plates and yeast paste. Two different embryo collection cages with *w*^*1118*^ and *βtub85Dtub85D_*cas9/+ males crossed to *w*^*1118*^ females served as a comparison control. Between four and six collections of 70-200 embryos each, were performed for each genotype. Every embryo collection was transferred in a separate fly vial and followed for over 36 h to count the number of embryos that did not hatch, the number of pupae, and female and male adults. Percent survival to each stage was calculated as the ratio between the number of individuals counted and the number of individuals expected for each genotype. The data for the two experiments and for each of the crosses are shown in Table 3 and Table 5.

## References

Buchman, A., and O. S. Akbari. 2019. “Site-specific transgenesis of the Drosophila melanogaster Y-chromosome using CRISPR/Cas9.” Insect Mol Biol 28 (1):65–73. doi: 10.1111/imb.12528.

Burt, A., and A. Deredec. 2018. “Self-limiting population genetic control with sex-linked genome editors.” Proc Biol Sci 285 (1883). doi: 10.1098/rspb.2018.0776.

Chan, Y. S., D. A. Naujoks, D. S. Huen, and S. Russell. 2011. “Insect population control by homing endonuclease-based gene drive: an evaluation in Drosophila melanogaster.” Genetics 188 (1):33–44 doi: 10.1534/genetics.111.127506.

Devlin, E. E., L. Dacosta, N. Mohandas, G. Elliott, and D. M. Bodine. 2010. “A transgenic mouse model demonstrates a dominant negative effect of a point mutation in the RPS19 gene associated with Diamond-Blackfan anemia.” Blood 116 (15):2826–35 doi: 10.1182/blood-2010-03-275776.

Dilthey, A. T., C. Jain, S. Koren, and A. M. Phillippy. 2019. “Strain-level metagenomic assignment and compositional estimation for long reads with MetaMaps.” Nat Commun 10 (1):3066. doi: 10.1038/s41467-019-10934-2.

Galizi, R., L. A. Doyle, M. Menichelli, F. Bernardini, A. Deredec, A. Burt, B. L. Stoddard, N. Windbichler, and A. Crisanti. 2014. “A synthetic sex ratio distortion system for the control of the human malaria mosquito.” Nat Commun 5:3977. doi: 10.1038/ncomms4977.

Galizi, R., A. Hammond, K. Kyrou, C. Taxiarchi, F. Bernardini, S. M. O’Loughlin, P. A. Papathanos, T. Nolan, N. Windbichler, and A. Crisanti. 2016. “A CRISPR-Cas9 sex-ratio distortion system for genetic control.” Sci Rep 6:31139. doi: 10.1038/srep31139.

Hall, A. B., S. Basu, X. Jiang, Y. Qi, V. A. Timoshevskiy, J. K. Biedler, M. V. Sharakhova, R. Elahi, M. A. Anderson, X. G. Chen, I. V. Sharakhov, Z. N. Adelman, and Z. Tu. 2015. “SEX DETERMINATION. A male-determining factor in the mosquito Aedes aegypti.” Science 348 (6240):1268–70 doi: 10.1126/science.aaa2850.

Hamilton, W. D. 1967. “Extraordinary sex ratios. A sex-ratio theory for sex linkage and inbreeding has new implications in cytogenetics and entomology.” Science 156 (3774):477–88

Kaufman, T. C. 2017. “A Short History and Description of.” Genetics 206 (2):665–689 doi: 10.1534/genetics.117.199950.

Kim, M., S. Ekhteraei-Tousi, J. Lewerentz, and J. Larsson. 2018. “The X-linked 1.688 Satellite in.” Genetics 208 (2):623–632 doi: 10.1534/genetics.117.300581.

Kondo, S., and R. Ueda. 2013. “Highly improved gene targeting by germline-specific Cas9 expression in Drosophila.” Genetics 195 (3):715–21 doi: 10.1534/genetics.113.156737.

Krzywinska, E., N. J. Dennison, G. J. Lycett, and J. Krzywinski. 2016. “A maleness gene in the malaria mosquito Anopheles gambiae.” Science 353 (6294):67–9 doi: 10.1126/science.aaf5605.

Kuhn, G. C., H. Küttler, O. Moreira-Filho, and J. S. Heslop-Harrison. 2012. “The 1.688 repetitive DNA of Drosophila: concerted evolution at different genomic scales and association with genes.” Mol Biol Evol 29 (1):7–11 doi: 10.1093/molbev/msr173.

Lindsley, D. L., and E. H. Grell. 1969. “Spermiogenesis without chromosomes in Drosophila melanogaster.” Genetics 61 (1):Suppl:69–78.

Lu, K. L., and Y. M. Yamashita. 2017. “Germ cell connectivity enhances cell death in response to DNA damage in the.” Elife 6. doi: 10.7554/eLife.27960.

Mali, P., L. Yang, K. M. Esvelt, J. Aach, M. Guell, J. E. DiCarlo, J. E. Norville, and G. M. Church. 2013. “RNA-guided human genome engineering via Cas9.” Science 339 (6121):823–6 doi: 10.1126/science.1232033.

Marygold, S. J., J. Roote, G. Reuter, A. Lambertsson, M. Ashburner, G. H. Millburn, P. M. Harrison, Z. Yu, N. Kenmochi, T. C. Kaufman, S. J. Leevers, and K. R. Cook. 2007. “The ribosomal protein genes and Minute loci of Drosophila melanogaster.” Genome Biol 8 (10):R216. doi: 10.1186/gb-2007-8-10-r216.

McKim, K. S., J. B. Dahmus, and R. S. Hawley. 1996. “Cloning of the Drosophila melanogaster meiotic recombination gene mei-218: a genetic and molecular analysis of interval 15E.” Genetics 144 (1):215–28

Meccariello, A., M. Salvemini, P. Primo, B. Hall, P. Koskinioti, M. Dalikova, A. Gravina, M. A. Gucciardino, F. Forlenza, M. E. Gregoriou, D. Ippolito, S. M. Monti, V. Petrella, M. M. Perrotta, S. Schmeing, A. Ruggiero, F. Scolari, E. Giordano, K. T. Tsoumani, F. Marec, N. Windbichler, K. P. Arunkumar, K. Bourtzis, K. D. Mathiopoulos, J. Ragoussis, L. Vitagliano, Z. Tu, P. A. Papathanos, M. D. Robinson, and G. Saccone. 2019. “Maleness-on-the-Y (MoY) orchestrates male sex determination in major agricultural fruit fly pests.” Science 365 (6460):1457–1460 doi: 10.1126/science.aax1318.

Menon, D. U., C. Coarfa, W. Xiao, P. H. Gunaratne, and V. H. Meller. 2014. “siRNAs from an X-linked satellite repeat promote X-chromosome recognition in Drosophila melanogaster.” Proc Natl Acad Sci U S A 111 (46):16460–5 doi: 10.1073/pnas.1410534111.

Mikhaylova, L. M., and D. I. Nurminsky. 2011. “Lack of global meiotic sex chromosome inactivation, and paucity of tissue-specific gene expression on the Drosophila X chromosome.” BMC Biol 9:29. doi: 10.1186/1741-7007-9-29.

Newton M.E., Wood R.L. and Southern D.I. 1976. “A cytogenetic analysis of meiotic drive in the mosquito Aedes aegypti.” Genetica 46:297–318.

Papathanos, P. A., and N. Windbichler. 2018. “Redkmer: An Assembly-Free Pipeline for the Identification of Abundant and Specific X-Chromosome Target Sequences for X-Shredding by CRISPR Endonucleases.” CRISPR J 1 (1):88–98 doi: 10.1089/crispr.2017.0012.

Pinello, L., M. C. Canver, M. D. Hoban, S. H. Orkin, D. B. Kohn, D. E. Bauer, and G. C. Yuan. 2016. “Analyzing CRISPR genome-editing experiments with CRISPResso.” Nat Biotechnol 34 (7):695–7 doi: 10.1038/nbt.3583.

Port, F., and S. L. Bullock. 2016. “Augmenting CRISPR applications in Drosophila with tRNA-flanked sgRNAs.” Nat Methods 13 (10):852–4 doi: 10.1038/nmeth.3972.

Port, F., H. M. Chen, T. Lee, and S. L. Bullock. 2014. “Optimized CRISPR/Cas tools for efficient germline and somatic genome engineering in Drosophila.” Proc Natl Acad Sci U S A 111 (29):E2967–76. doi: 10.1073/pnas.1405500111.

Romeijn, R. J., M. M. Gorski, M. A. van Schie, J. N. Noordermeer, L. H. Mullenders, W. Ferro, and A. Pastink. 2005. “Lig4 and rad54 are required for repair of DNA double-strand breaks induced by P-element excision in Drosophila.” Genetics 169 (2):795–806 doi: 10.1534/genetics.104.033464.

Staeva-Vieira, E., S. Yoo, and R. Lehmann. 2003. “An essential role of DmRad51/SpnA in DNA repair and meiotic checkpoint control.” EMBO J 22 (21):5863–74 doi: 10.1093/emboj/cdg564.

Stewart, M. J., and R. Denell. 1993. “The Drosophila ribosomal protein S6 gene includes a 3’ triplication that arose by unequal crossing-over.” Mol Biol Evol 10 (5):1041–7 doi: 10.1093/oxfordjournals.molbev.a040053.

Sweeny, T L, and A R Barr. 1978. “Sex Ratio Distortion Caused by Meiotic Drive in a Mosquito, Culex pipiens L.” Genetics 88 (3):427–446

Vibranovski, M. D., Y. E. Zhang, C. Kemkemer, H. F. Lopes, T. L. Karr, and M. Long. 2012. “Re-analysis of the larval testis data on meiotic sex chromosome inactivation revealed evidence for tissue-specific gene expression related to the drosophila X chromosome.” BMC Biol 10:49; author reply 50. doi: 10.1186/1741-7007-10-49.

White-Cooper, Helen. 2012. “Tissue, cell type and stage-specific ectopic gene expression and RNAi induction in the Drosophila testis.” Spermatogenesis 2 (1):11–22 doi: 10.4161/spmg.19088.

Windbichler, N., P. A. Papathanos, and A. Crisanti. 2008. “Targeting the X chromosome during spermatogenesis induces Y chromosome transmission ratio distortion and early dominant embryo lethality in Anopheles gambiae.” PLoS Genet 4 (12):e1000291. doi: 10.1371/journal.pgen.1000291.

