## Supplementary Data 1 for "A fly model establishes distinct mechanisms for synthetic CRISPR/Cas9 sex distorters"

**Table 1.** The F1 progeny from the crosses between *βtub85D_cas9/sgRNA* and wild type *w* lines

| female X male | females | males | females % | males % |
|---|---|---|---|---|
| *w* X ***esi-2. 1_1***/*βtub85D-cas9*$^{\#4}$ | 147 | 176 | 45.5 | 54.5 |
| ***esi-2.1_1***/*βtub85D-cas9*$^{\#4}$ X *w* | 180 | 148 | 54.9 | 45.1 |
| *w* X ***CG33235_1***/*βtub85D-cas9*$^{\#4}$ | 174 | 215 | 44.7 | 55.3 |
| ***CG33235_1***/*βtub85D-cas9*$^{\#4}$ X *w* | 151 | 148 | 50.5 | 49.5 |
| *w* X ***Muc14a_1***/*βtub85D-cas9*$^{20F}$ | 1776 | 1711 | 51.1 | 48.9 |
| *w* X ***Muc14a_2***/*βtub85D-cas9*$^{20F}$ | 1472 | 1623 | 47.7 | 52.3 |
| *w* X ***Muc14a_3***/*βtub85D-cas9*$^{20F}$ | 1893 | 1800 | 51.4 | 48.6 |
| *w* X ***Muc14a_4***/*βtub85D-cas9*$^{20F}$ | 1707 | 1620 | 51.2 | 48.8 |
| ***Muc14a_5***/*βtub85D-cas9*$^{\#4}$X *w* | 249 | 238 | 51.1 | 48.9 |
| *w* X ***Muc14a_5***/*βtub85D-cas9*$^{\#4}$ | 271 | 274 | 49.7 | 50.3 |
| *w* X ***Muc14a_6***/*βtub85D-cas9*$^{20F}$ | 1199 | 1895 | 38.5 | 61.5 |
| *w* X *βtub85D-cas9*$^{20F}$/ + | 2387 | 2183 | 52.5 | 47.5 |
| *w* X ***CG15040_1***/*βtub85D-cas9*$^{\#4}$ | 210 | 180 | 53.8 | 46.2 |
| ***CG15040_1***/*βtub85D-cas9*$^{\#4}$ X *w* | 196 | 197 | 49.9 | 50.1 |
| *w* X ***hydra_1***/*βtub85D-cas9*$^{20F}$ | 335 | 309 | 52.0 | 48.0 |
| ***hydra_1***/*βtub85D-cas9*$^{20F}$ X *w* | 350 | 294 | 54.3 | 45.7 |
| *w* X ***RpS6_1***/*βtub85D-cas9*$^{20F}$ | 1489 | 1481 | 50.0 | 50.0 |
| *w* X ***RpS6_2***/*βtub85D-cas9*$^{20F}$ | 170 | 2659 | 6.2 | 93.8 |
| *w* X ***RpS5a_1***/*βtub85D-cas9*$^{20F}$ | 1017 | 1325 | 43.4 | 56.6 |
| *w* X ***RpS5a_2***/*βtub85D-cas9*$^{20F}$ | 1610 | 1595 | 50.3 | 49.7 |
| *w* X*βtub85D-cas9*$^{20F}$/ + | 1066 | 1114 | 49.2 | 50.8 |

**Table 2.** The F1 progeny of crosses with one and two *βtub85D_cas9* and *nos-cas9*

| | female X male | females | males | females % (STDEV) | males % (STDEV) | *w* eye females |
|---|---|---|---|---|---|---|
| **1 X *βtub85D-cas9*** | *w* X ***Muc14a_3***/+; ***Muc14a_4***/*βtub85D-cas9*$^{\#4}$ | 201 | 230 | 46.6 | 53.4 | |
| | ***Muc14a_3***/+; ***Muc14a_4***/*βtub85D-cas9*$^{\#4}$ X *w* | 185 | 192 | 49.1 | 50.9 | |
| | *w* X ***Muc14a_6***/+; ***hydra_1***/*βtub85D-cas9*$^{20F}$ | 1106 | 1558 | 41.0 (5.2) | 59.0 (5.2) | |
| | *w* X *βtub85D-cas920F/+* | 2387 | 2183 | 52.5 (5.0) | 47.5 (5.0) | |
| | *w* X *w+*; ***gRNA array****/*βtub85D-cas9*$^{20F}$ | 179 | 197 | 47.6 | 52.4 | 2.8 |
| | *w/w+*; ***gRNA array****/*βtub85D-cas9*$^{20F}$ X *w* | 213 | 259 | 45.1 | 54.9 | 64.8 |
| | *w* X ***Muc14a_6***/*βtub85D-cas9*$^{\#10\text{-}1}$ | 78 | 118 | 39.8 | 60.2 | |
| **2 X *βtub85D-cas9*** | *w* X *βtub85D-cas9*$^{\#15\text{-}1}$/+; *βtub85D-cas9*$^{\#10\text{-}1}$/***Muc14a_6*** | 319 | 273 | 53.9 | 46.1 | |
| | *w* X *βtub85D-cas9*$^{\#15\text{-}1}$/+; *βtub85D-cas9*$^{\#10\text{-}1}$/***esi2.1_1*** | 207 | 204 | 50.4 | 49.6 | |
| | *βtub85D-cas9*$^{\#15\text{-}1}$/+; *βtub85D-cas9*$^{\#10\text{-}1}$/***esi2.1_1*** X *w* | 146 | 152 | 49.0 | 51.0 | |
| ***nos-cas9*** | *w* X ***Muc14a_6***/*nos_cas9* | 360 | 354 | 50.4 | 49.6 | |
| | *w* X *nos_cas9/+* | 744 | 736 | 49.7 (3.5) | 50.3 (3.5) | |

**Muc14a_3, hydra_1, esi-2.1_1, w_ex3-2*

**Table 3.** Determination of survival in the F1 progeny of *Muc14a_6/ βtub85D_cas9* individuals

| Males X *w* | Vial # | T ($^0$C) | # of embryos | hatched embryos | # pupae | adults | females | males | % hatched embryos | % pupae | % adults | % females | % males |
|---|---|---|---|---|---|---|---|---|---|---|---|---|---|
| *Muc14a_6/βtub85D-cas9*$^{20F}$ | 1 | 25 | 140 | 82 | 77 | 77 | 26 | 51 | 58.6 | 55.0 | 55.0 | 34.0 | 66.0 |
| | 2 | 25 | 158 | 83 | 82 | 79 | 30 | 49 | 52.5 | 51.9 | 50.0 | 38.0 | 62.0 |
| | 3 | 25 | 154 | 100 | 96 | 93 | 28 | 65 | 64.9 | 62.3 | 60.4 | 30.0 | 70.0 |
| | 4 | 25 | 50 | 37 | 31 | 31 | 10 | 21 | 74.0 | 62.0 | 62.0 | 32.0 | 68.0 |
| | 5 | 25 | 175 | 137 | 115 | 111 | 37 | 74 | 78.3 | 65.7 | 63.4 | 33.0 | 67.0 |
| | 6 | 25 | 146 | 96 | 91 | 88 | 30 | 58 | 65.8 | 62.3 | 60.3 | 34.0 | 66.0 |
| Average | | | | | | | | | 65.7 | 59.9 | 58.5 | 33.6 | 66.4 |
| STDEV | | | | | | | | | 9.5 | 5.3 | 5.1 | 2.6 | 2.6 |
| *βtub85D-cas9*$^{20F}$*/+* | 1 | 25 | 125 | 71 | 68 | 68 | 31 | 37 | 56.8 | 54.4 | 54.4 | 45.6 | 54.4 |
| | 2 | 25 | 151 | 103 | 103 | 99 | 44 | 55 | 68.2 | 68.2 | 65.6 | 44.4 | 55.6 |
| | 3 | 25 | 161 | 119 | 83 | 79 | 45 | 34 | 73.9 | 51.6 | 49.1 | 57.0 | 43.0 |
| | 4 | 25 | 152 | 97 | 104 | 104 | 52 | 52 | 63.8 | 68.4 | 68.4 | 50.0 | 50.0 |
| | 5 | 25 | 70 | 56 | 48 | 47 | 33 | 15 | 80.0 | 68.6 | 67.1 | 70.2 | 31.9 |
| Average | | | | | | | | | 68.5 | 62.2 | 60.9 | 53.4 | 47.0 |
| STDEV | | | | | | | | | 9.0 | 8.5 | 8.7 | 10.6 | 9.8 |
| *w* | 1 | 25 | 161 | 154 | 152 | 150 | 72 | 78 | 95.7 | 94.4 | 93.2 | 48.0 | 52.0 |
| | 2 | 25 | 184 | 175 | 159 | 159 | 98 | 61 | 95.1 | 86.4 | 86.4 | 61.6 | 38.4 |
| | 3 | 25 | 161 | 153 | 144 | 143 | 62 | 81 | 95.0 | 89.4 | 88.8 | 43.4 | 56.6 |
| | 4 | 25 | 150 | 132 | 119 | 118 | 58 | 60 | 88.0 | 79.3 | 78.7 | 49.2 | 50.8 |
| Average | | | | | | | | | 93.4 | 87.4 | 86.8 | 50.5 | 49.5 |
| STDEV | | | | | | | | | 3.6 | 6.3 | 6.1 | 7.8 | 7.8 |

**Table 4.** The F1 progeny from the crosses between *Muc14a_6* gRNA + mutant; *βtub85D_cas9* and wild type *w* lines

| | | Wild type *w* mothers | | | |
|---|---|---|---|---|---|
| | | **females** | **males** | **females % (STDEV)** | **males % (STDEV)** |
| **Fathers** | *βtub85D_cas9*$^{20F}$/ + | 1257 | 1266 | 49.8 (2.4) | 53.6 (2.4) |
| | ***Muc14a_6***/ *βtub85D_cas9*$^{20F}$ | 1199 | 1895 | 38.5 (4.8) | 61.5 (4.8) |
| | ***DNAlig4***$^{\#57}$; ***Muc14a_6***/ *βtub85D_cas9*$^{20F}$ | 1404 | 3043 | 31.7 (4.4) | 68.3 (4.4) |
| | *βtub85D_cas9*$^{20G}$/+ | 1273 | 1319 | 49.0 (3.7) | 51.0 (3.7) |
| | ***Muc14a_6***/ *βtub85D_cas9*$^{20G}$ | 1044 | 1607 | 39.5 (4.3) | 60.5 (4.3) |
| | ***Muc14a_6***/ *βtub85D_cas9*$^{20G}$; ***Spn-A***$^{057}$/ ***Df(3R)X3F*** | 870 | 1518 | 35.8 (7.2) | 64.2 (7.2) |
| | ***Muc14a_6***/ *βtub85D_cas9*$^{20G}$; ***Spn-A***$^{093}$/ ***Df(3R)X3F*** | 850 | 1257 | 40.3 (3.0) | 59.7 (3.0) |
| | ***Muc14a_6/+;*** *βtub85D_cas9*$^{20F}$/ + | 1089 | 1865 | 36.4 (7.6) | 63.6 (7.6) |
| | ***nos_GAL4*** *βtub85D_cas9*$^{20F}$/ + | 1100 | 985 | 52.7 (3.5) | 47.3 (3.5) |
| | ***UAS_spectrinRNAi/+; Muc14a_6/ nos_GAL4*** *βtub85D_cas9*$^{20F}$ | 774 | 1241 | 38.6 (2.9) | 61.4 (2.9) |

**Table 5.** The F1 progeny from the crosses between *βtub85D_cas9* combined with ribosomal genes sgRNAs and wild type *w* females

| | | Wild type *w* mothers | | | | | |
|---|---|---|---|---|---|---|---|
| | | # of crosses | females | males | males % (STDEV) | Average F1 # per single cross | F1 mutant to control relative ratio |
| **Fathers** | *βtub85D_cas9$^{20F}$/ +* | 9 | 1066 | 1114 | 50.8 (2.9) | 214.4 (55) | 1 |
| | ***RpS5a_1**/βtub85D_cas9$^{20F}$* | 15 | 1017 | 1325 | 56.6 (5.9) | 156.1 (21.5) | 0.73 |
| | ***RpS5a_2**/βtub85D_cas9$^{20F}$* | 15 | 1610 | 1595 | 49.7 (4.0) | 213.7 (30.8) | 0.99 |
| | ***RpS6_1**/βtub85D_cas9$^{20F}$* | 14 | 1489 | 1481 | 50.0 (4.4) | 212.0 (25.3) | 0.99 |
| | ***RpS6_2**/βtub85D_cas9$^{20F}$* | 19 | 170 | 2659 | 93.8 (3.4) | 175.5 (38.9) | 0.82 |
| | ***RpS6_2 RpS5a_1**/βtub85D_cas9$^{20F}$* | 32 | 522 | 3848 | 88.1 (5.1) | 136.6 (45.4) | 0.63 |
| | ***RpS6_2 RpS6_1**/βtub85D_cas9$^{20F}$* | 24 | 493 | 3236 | 87.0 (4.3) | 155.4 (38.8) | 0.72 |
| | ***Muc14a_6**/+;**RpS6_2**/βtub85D_cas9$^{20F}$* | 12 | 63 | 1477 | 95.8 (4.2) | 128.3 (63.0) | 0.60 |

**Table 6.** Determination of survival in the F1 progeny of *RpS6_2/ βtub85Dcas9* individuals

| Males X *w* | Vial # | T ($^0$C) | # of embryos | hatched embryos | # pupae | adults | females | males | % hatched embryos | % pupae | % adults | % females | % males |
|---|---|---|---|---|---|---|---|---|---|---|---|---|---|
| *βtub85Dcas9/**RpS6_2*** | 1 | 25 | 56 | 46 | 45 | 40 | 2 | 38 | 82.1 | 80.4 | 71.4 | 3.6 | 67.9 |
| | 2 | | 142 | 114 | 89 | 63 | 7 | 56 | 80.3 | 62.7 | 44.4 | 4.9 | 39.4 |
| | 3 | | 176 | 131 | 108 | 83 | 7 | 76 | 74.4 | 61.4 | 47.2 | 4.0 | 43.2 |
| | 4 | | 154 | 93 | 82 | 63 | 7 | 56 | 60.4 | 53.2 | 40.9 | 4.5 | 36.4 |
| | 5 | | 175 | 130 | 115 | 91 | 6 | 85 | 74.3 | 65.7 | 52.0 | 3.4 | 48.6 |
| | 6 | | 64 | 41 | 30 | 22 | 4 | 18 | 64.1 | 46.9 | 34.4 | 6.3 | 28.1 |
| Average | | | | | | | | | 72.6 | 61.7 | 48.4 | 4.5 | 43.9 |
| STDEV | | | | | | | | | 8.7 | 11.5 | 12.8 | 1.1 | 13.6 |
| *βtub85D_cas9/+* | 1 | 25 | 137 | 121 | 108 | 104 | 50 | 54 | 88.3 | 78.8 | 75.9 | 36.5 | 39.4 |
| | 2 | | 135 | 126 | 113 | 108 | 59 | 49 | 93.3 | 83.7 | 80.0 | 43.7 | 36.3 |
| | 3 | | 68 | 59 | 57 | 56 | 25 | 31 | 86.8 | 83.8 | 82.4 | 36.8 | 45.6 |
| | 4 | | 137 | 118 | 113 | 105 | 62 | 43 | 86.1 | 82.5 | 76.6 | 45.3 | 31.4 |
| | 5 | | 127 | 108 | 117 | 91 | 40 | 51 | 85.0 | 92.1 | 71.7 | 31.5 | 40.2 |
| Average | | | | | | | | | 87.9 | 84.2 | 77.3 | 38.7 | 38.6 |
| STDEV | | | | | | | | | 3.3 | 4.9 | 4.1 | 5.7 | 5.2 |
| *w* | 1 | 25 | 135 | 133 | 126 | 123 | 64 | 59 | 98.5 | 93.3 | 91.1 | 47.4 | 44.4 |
| | 2 | | 50 | 48 | 45 | 44 | 24 | 20 | 96.0 | 90.0 | 88.0 | 48.0 | 41.7 |
| | 3 | | 169 | 162 | 145 | 139 | 68 | 71 | 95.9 | 85.8 | 82.2 | 40.2 | 43.8 |
| | 4 | | 144 | 133 | 128 | 115 | 50 | 65 | 92.4 | 88.9 | 79.9 | 34.7 | 48.9 |
| | 5 | | 164 | 144 | 135 | 127 | 57 | 70 | 87.8 | 82.3 | 77.4 | 34.8 | 48.6 |
| Average | | | | | | | | | 94.1 | 88.1 | 83.7 | 41.0 | 45.5 |
| STDEV | | | | | | | | | 4.1 | 4.2 | 5.7 | 6.5 | 3.2 |

**Table S1.** Testing the efficiencies of attP and piggyBac integrated *cas9* and *cpf1* lines

| | | Wild type *w* mothers | | | | |
|---|---|---|---|---|---|---|
| | | | | Females | | |
| | **Integration** | | **T ($^0$C)** | ***$w^+$*** | ***w*** | **% w** |
| **Fathers** | **attP** | *$w^+$; **βtub85D-cas9$^{\#15-1}$**/+; w_ex3-2 #2/+* | 25 | 338 | 54 | 13.8 |
| | | *$w^+$; **βtub85D-cas9$^{\#10-1}$**/w_ex3-2 #2* | 25 | 59 | 16 | 21.3 |
| | | *$w^+$; **βtub85D-cas9$^{\#4}$**/+; w_ex3-2 #2* | 25 | 62 | 18 | 18.2 |
| | | *$w^+$; **βtub85D-cas9$^{\#15-1}$**; **βtub85D-Cas9$^{\#10-1}$**/w_ex3-2 #2* | 25 | 55 | 67 | 54.9 |
| | | *$w^+$; **βtub85D-LbCpf1$^{\#2}$**/w_ex3-1 #2* | 29 | 175 | 1 | 0.6 |
| | **piggyBac** | *$w^+$ **βtub85D-cas9$^{\#20D}$**; w_ex3-2 #2/+* | 25 | 1204 | 429 | 25.5 |
| | | *$w^+$; **βtub85D-cas9$^{\#20G}$**/+; w_ex3-2 #2/+* | 25 | 109 | 62 | 36.3 |
| | | *$w^+$; **βtub85D-cas9$^{\#20F}$**/w_ex3-2 #2* | 25 | 155 | 169 | 52.2 |
| | | *$w^+$; **βtub85D-cas9$^{\#20C}$**/w_ex3-2 #2* | 25 | 248 | 123 | 33.2 |
| | | *$w^+$; **βtub85D-cas9$^{\#20F}$**/Array1_1* | 25 | 164 | 5 | 2.8 |
| | **attP** | *$w^+$; **nos-cas9**/w_ex3-2 #2* | 25 | 6 | 177 | 96.7 |

| **Array_1** |
|---|
| *Muc14a_4* |
| *hydra_1* |
| *esi-2.1_1* |
| *w_ex3-2* |
