## Supplementary material for "A fly model establishes distinct mechanisms for synthetic CRISPR/Cas9 sex distorters": Table S1

**Primers used in this study**. The sequence of the gRNA in each oligonucleotide is underlined

| Primer name | Description | Primer sequence |
|---|---|---|
| βtub85D_F | pYSC47w+_βtub85D_cas9_βtub56D | ATACTTATTAGTCAGTCCAGAAACAACTTTGGCACCTGCAGTCAATATCAATCGTATCATCTGGTCGG |
| βtub85D-Cas9_R | pYSC47w+_βtub85D_cas9_βtub56D | TCGTGGTCCTTATAGTCCATTTTGATAGTAAAGTTAGGGCCCCTTTTT |
| βtub85D-Cas9_F | pYSC47w+_βtub85D_cas9_βtub56D | GCCCTAACTTTACTATCAAAATGGACTATAAGGACCACGACGG |
| Cas9-βtub56D-3'UTR_R | pYSC47w+_βtub85D_cas9_βtub56D | TTTAGTTCTCGTCGACCTCATTACTTTTTCTTTTTTGCCTGGCCG |
| Casβtub56_F | pYSC47w+_βtub85D_cas9_βtub56D | AGGCAAAAAAGAAAAAGTAATGAGGTCGACGAGAACTAAATTCG |
| βtub56D_R | pYSC47w+_βtub85D_cas9_βtub56D | GTGGACACGCTAGACCAAATGTGTTCTGTGATGACCTGCATCATGTCCTCTTCAAGGGCGAA |
| βtub85D pYSC47_F | pYSC47w+_βtub85D_Lbcpf1_βtub56D | ATACTTATTAGTCAGTCCAGAAACAACTTTGGCACCTGCAGTCAATATCAAT |
| βtub85DhLbCpf1_R | pYSC47w+_βtub85D_Lbcpf1_βtub56D | AACTTCTCCAGCTTGCTCATTTTGATAGTAAAGTTAGGGCCCCTTTTT |
| βtub85DhLbCpf1_F | pYSC47w+_βtub85D_Lbcpf1_βtub56D | GCCCTAACTTTACTATCAAAATGAGCAAGCTGGAGAAGTTTACAAAC |
| hLbCpf1βtub56D_R | pYSC47w+_βtub85D_Lbcpf1_βtub56D | TTTAGTTCTCGTCGACCTCATTAGGCATAGTCGGGGACATCATATGG |
| hLbCpf1βtub56D_F | pYSC47w+_βtub85D_Lbcpf1_βtub56D | ATGTCCCCGACTATGCCTAATGAGGTCGACGAGAACTAAATTCGAATCG |
| βtub56DpYSC47_R | pYSC47w+_βtub85D_Lbcpf1_βtub56D | GTGGACACGCTAGACCAAATGTGTTCTGTGATGACCTGCATCATGTCCTCTT |
| 58537687_F | in vitro Cas9 test | AAAATCTCCACTCCAGTTGAGTACAACTTCC |
| Muc14_F | in vitro Cas9 test | GTCGGTTCTGTTGTTAATTGAGGTA |
| Cloning Analysis Forward Primer | in vitro Cas9 test | ACCTGCCAACCAAAGCGAGAAC |
| Cloning Analysis Reverse Primer | in vitro Cas9 test | TCAGGGTTATTGTCTCATGAGCG |
| gRNA-Muc14a_6_F | cloning into pCFD3 | GTCGGAACAGCTCAAGAGGAGACAT |
| gRNA-Muc14a_6_R | cloning into pCFD3 | AAACATGTCTCCTCTTGAGCTGTTC |
| gRNA-Muc14a_4_F | cloning into pCFD3 | GTCGGTTCTGTTGTTAATTGAGGTA |
| gRNA-Muc14a_4_R | cloning into pCFD3 | AAACTACCTCAATTAACAACAGAAC |
| gRNA-Muc14a_5_F | cloning into pCFD3 | GTCGGAGAAGAGCAAACAGCTCAAG |
| gRNA-Muc14a_5_R | cloning into pCFD3 | AAACCTTGAGCTGTTTGCTCTTCTC |
| gRNA-Muc14a_3_F | cloning into pCFD3 | GTCGGCTTTCTTCTGTTGTTAATTG |
| gRNA-Muc14a_3_R | cloning into pCFD3 | AAACCAATTAACAACAGAAGAAAGC |
| gRNA-hydra_1_F | cloning into pCFD3 | GTCGGACAAATTTTTGATTTCAGAG |
| gRNA-hydra_1_R | cloning into pCFD3 | AAACCTCTGAAATCAAAAATTTGTC |
| gRNA-esi-2.1_1_F | cloning into pCFD3 | GTCGGACTTGTTTGAGTCCAACTAC |
| gRNA-esi-2.1_1_R | cloning into pCFD3 | AAACGTAGTTGGACTCAAACAAGTC |
| gRNA-CG33235_1_F | cloning into pCFD3 | GTCGGTCATATCGGGGCCTCCTTGC |
| gRNA-CG33235_1_R | cloning into pCFD3 | AAACGCAAGGAGGCCCCGATATGAC |
| gRNA-CG15040_F | cloning into pCFD3 | GTCGGTGTTGTTCTGATTCTTGCTT |
| gRNA-CG15040_R | cloning into pCFD3 | AAACAAGCAAGAATCAGAACAACAC |
| gRNA-Muc14a_1_F | cloning into pCFD3 | GTCGGATCCATCGTGCGAAGCCAA |
| gRNA-Muc14a_1_R | cloning into pCFD3 | AAACTTGGCTTCGCACGATGGATC |
| gRNA-Muc14a_2_F | cloning into pCFD3 | GTCGGAGTAAAGACCTCACAACTA |
| gRNA-Muc14a_2_R | cloning into pCFD3 | AAACTAGTTGTGAGGTCTTTACTC |
| gRNA-w-ex3-2_F | cloning into pCFD3 | GTCGGGTGATGGGCAGTTCCGGTGC |
| gRNA-w-ex3-2_R | cloning into pCFD3 | AAACGCACCGGAACTGCCCATCACC |
| gRNA-w-ex3-1_F | cloning into pCFD3 w/o scaffold | GTCGGAATTTCTACTAAGTGTAGATGCCGTGATGGGCAGTTCCGG |
| gRNA-w-ex3-1_R | cloning into pCFD3 w/o scaffold | AAAACCGGAACTGCCCATCACGGCATCTACACTTAGTAGAAATTC |
| gRNA-RpS5a_1_F | cloning into pCFD3 | GTCGGGAGACCTTCGAGGAGCCAG |
| gRNA-RpS5a_1_R | cloning into pCFD3 | AAACCTGGCTCCTCGAAGGTCTCC |
| gRNA-RpS5a_2_F | cloning into pCFD3 | GTCGGCGAAGTTGCTGAAAACGTGG |
| gRNA-RpS5a_2_R | cloning into pCFD3 | AAACCCACGTTTTCAGCAACTTCGC |
| gRNA-RpS6_1_F | cloning into pCFD3 | GTCGGTCGCATTGCAAACTTACCGT |
| gRNA-RpS6_1_R | cloning into pCFD3 | AAACACGGTAAGTTTGCAATGCGAC |
| gRNA-RpS6_2_F | cloning into pCFD3 | GTCGGCCGGCGGCAACGACAAGCA |
| gRNA-RpS6_2_R | cloning into pCFD3 | AAACTGCTTGTCGTTGCCGCCGGC |
| RpS6_2 pCFD4_F | Multiplex cloning into pCFD4 | TATATATAGGAAAGATATCCGGGTGAACTTCGCCGGCGGCAACGACAAGCAGTTTTAGAGCTAGAAATAGCAAG |
| RpS6_1 pCFD4_R | Multiplex cloning into pCFD4 | ATTTTAACTTGCTATTTCTAGCTCTAAAACACGGTAAGTTTGCAATGCGACGACGTTAAATTGAAAATAGGTC |
| RpS5a_1 pCFD4_R | Multiplex cloning into pCFD4 | ATTTTAACTTGCTATTTCTAGCTCTAAAACCTGGCTCCTCGAAGGTCTCCGACGTTAAATTGAAAATAGGTC |
| Array1_Cas9_PCR1_F | Multiplex cloning into pCFD5 | GCGGCCCGGGTTCGATTCCCGGCCGATGCATTCTGTTGTTAATTGAGGTAGTTTTAGAGCTAGAAATAGCAAG |
| Array1_Cas9_PCR1_R | Multiplex cloning into pCFD5 | CTCTGAAATCAAAAATTTGTTGCACCAGCCGGGAATCGAACCC |
| Array1_Cas9_PCR2_F | Multiplex cloning into pCFD5 | ACAAATTTTTGATTTCAGAGGTTTTAGAGCTAGAAATAGCAAG |
| Array1_Cas9_PCR2_R | Multiplex cloning into pCFD5 | GTAGTTGGACTCAAACAAGTTGCACCAGCCGGGAATCGAACCC |
| Array1_Cas9_PCR3_F | Multiplex cloning into pCFD5 | ACTTGTTTGAGTCCAACTACGTTTTAGAGCTAGAAATAGCAAG |
| Array1_Cas9_PCR3_R | Multiplex cloning into pCFD5 | ATTTTAACTTGCTATTTCTAGCTCTAAAACGCACCGGAACTGCCCATCACCTGCACCAGCCGGGAATCGAACCC |
| repeat_56910823_ illumina_F | Muc14a amplicon sequencing | ACACTCTTTCCCTACACGACGCTCTTCCGATCTTCAAGAAGATACAAGGACAC |
| repeat_Muc14a_3_illumina_R | Muc14a amplicon sequencing | GACTGGAGTTCAGACGTGTGCTCTTCCGATCTCTTTCTTCTGTTGTTAATTGAGG |
| RpS6_1F | RpS6 amplicon sequencing | CCAAAAGCTATTCGAAGTGGTC |
| RpS6_1R | RpS6 amplicon sequencing | CACTCTTTCAAGGCGTACAAAA |
